# 2’-O-Methyl-guanosine 3-base RNA fragments mediate essential natural TLR7/8 antagonism

**DOI:** 10.1101/2024.07.25.605091

**Authors:** Arwaf S. Alharbi, Sunil Sapkota, Zhikuan Zhang, Ruitao Jin, W. Samantha N. Jayasekara, Erandi Rupasinghe, Mary Speir, Lorna Wilkinson-White, Roland Gamsjaeger, Liza Cubeddu, Julia I. Ellyard, Daniel S. Wenholz, Alexandra L. McAllan, Refaya Rezwan, Le Ying, Hani Hosseini Far, Josiah Bones, Sitong He, Dingyi Yu, Kim A. Lennox, Paul J. Hertzog, Carola G. Vinuesa, Mark A. Behlke, Umeharu Ohto, Olivier F. Laczka, Ben Corry, Toshiyuki Shimizu, Michael P. Gantier

## Abstract

Recognition of RNA fragments by Toll-like receptors (TLR) 7 and 8 is a key contributor to the initiation of a protective innate immune response against pathogens. A long-standing enigma is how degradation products of host RNAs, generated by the daily phagocytic clearance of billions of apoptotic cells, fail to activate TLR7 and TLR8 signalling^1^. Here, we report that select 2’-O-methyl (2’-Ome) guanosine RNA fragments as short as 3 bases, including those derived from host-RNAs, are potent TLR7 and TLR8 antagonists that reduce TLR7 sensing *in vivo*. Mechanistically, antagonistic fragments are directed towards a distinct binding site on these proteins by 5’-end 2’-Ome guanosine. Our results indicate that host-RNAs evade detection by TLR7/8 due to a pool of abundant host ribosomal 2’-Ome-modified RNA fragments that naturally antagonize TLR7 and TLR8 sensing to avoid auto-immunity. Crucially, rare TLR7 and TLR8 mutations located at this antagonistic site decrease the inhibitory activity of 2’-Ome guanosine RNA fragments and lead to auto-immunity in patients. Our findings also establish that select chemically synthesised 3-base oligonucleotides can harness the protective anti-inflammatory activity of this natural immune checkpoint for therapeutic targeting of TLR7-driven diseases.

**One Sentence Summary:** Short 2’-O-Methyl RNA fragments are natural TLR7/8 antagonists

---

Chromosome X-encoded endosomal Toll-like receptors (TLR) 7 and 8 play critical roles in the initiation of innate immune responses to pathogens through the detection of phagocytosed RNAs. Consequently, male patients deficient in TLR7 are more prone to life-threatening SARS-CoV-2 infections^2^. These receptors normally evade detection of phagocytosed host-RNAs, however rare mutations in TLR7 and TLR8 alter this evasion and promote autoimmunity in patients ^3–6^. Critically, how this differentiation between pathogenic and host RNAs is achieved has remained unclear to date. While initially described as innate immune sensors specialised in the detection of single-stranded RNAs (ssRNAs) as short as ∼20 bases^7–9^, recent structural studies have demonstrated that TLR7 and TLR8 dimers in fact bind to RNA degradation products through the engagement of two distinct ligand binding sites^10–13^.

The first binding site, hereafter referred to as site 1, is highly conserved between the two TLRs and binds the ultimate RNA degradation products in the form of single uridine residues for TLR8 and single guanosine (or guanosine 2’,3’-cyclic phosphate [2’,3’-cGMP]) residues for TLR7^11–14^. Site 1 can also bind to base analogues and imidazoquinolines, such as the FDA-approved imiquimod (TLR7), resiquimod/R848 (TLR7 and TLR8), or their derivatives, and its engagement is sufficient to promote functional activation of both receptors^10–13^. The second site (site 2), has been shown to bind short uridine-containing RNA motifs of 2-3 bases, such as UUU and UG, which facilitate guanosine and uridine binding to site 1 of TLR7 and TLR8, respectively^11,12^. As such, engagement of site 2 synergises with that of site 1 to strongly potentiate TLR7 and TLR8 sensing^11–14^. RNAse T2 and RNAse 2 are essential for processing of ssRNA into short fragments for TLR7 and TLR8 activation, as their absence blunts TLR7 and TLR8 activation by ssRNA and bacterial RNA^15–17^.

Following the initial description of RNA sensing by TLR7 and TLR8, inhibition by modifications in mammalian RNA was reported. As such, several studies have shown that sugar and base modification of uridines, guanosines, and adenosines can blunt TLR7 and TLR8 sensing of otherwise potent TLR7/8 agonistic ssRNAs^18–22^. Methylation of the ribose 2’-group of the RNA bases, known as 2’-O-methyl (2’-OMe) modification, was proposed to generate antagonistic ssRNAs with a stronger affinity to TLR7 *in vitro*, blocking sensing of otherwise activating ssRNAs^20,22,23^. Given that >100 residues are known to be 2’-OMe-modified in mammalian ribosomal RNA^24–26^, representing ∼80% of mammalian cellular RNA, it was inferred that such modification of endogenous RNAs was directly involved in limiting TLR7 and TLR8 activation by phagocytosed endogenous RNA^18^. This concept was further supported by reports that select bacterial transfer RNAs (tRNA), containing a 2’-OMe motif (*e.g.*, Gm18), could block TLR7 recognition of RNA derived from specific bacterial strains^27,28^. Notably, exactly how 2’-OMe-modified RNA antagonised RNA sensing, mechanistically, was not investigated in these studies – the dogma being that abundant endogenous RNAs contained sporadic modifications sufficient to tag them as self.

In this context, the recent discovery that TLR7 and TLR8 in fact detect very short RNA degradation fragments that, given the low frequency of RNA-modifications within longer RNAs, are mostly unmodified, raises the question of how phagocytosed fragments of self-RNAs evade aberrant recognition in homeostasis. Here we reveal that 2’-OMe-modified oligonucleotides as short as 3-bases can bind to TLR7 and TLR8 to modulate their function in a sequence-specific manner. Our systematic sequence analyses together with cryo-electron microscopy (cryo-EM) structural studies of *Macaca mulatta* TLR7 with 3-base oligonucleotides and molecular dynamics simulation established an essential role for 2’-OMe guanosine in TLR7 and TLR8 antagonism. This occurs at a separate binding site, distinct from the familiar site 1 and site 2. Notably, naturally occurring rare mutations of the TLR7 and TLR8 residues underpinning antagonism by 2’-OMe guanosine RNA fragments leads to autoimmunity in patients, establishing the importance of this interaction for homeostasis. Taken together our findings suggest that abundant 2’-OMe guanosine motifs found in ribosomal RNA and its degradation products, act as *bona fide* antagonists of TLR7 and TLR8, helping to safeguard against aberrant sensing of self-RNA. These results also reveal the therapeutic potential of very short synthetic 2’-OMe guanosine-RNA fragments to curb aberrant TLR7/8-driven inflammation.

## Results

### 3-base long 2’-OMe-modified oligonucleotides modulate TLR8 sensing

Spontaneously endocytosed DNA and 2’-OMe-modified phosphorothioate (PS) oligonucleotides (oligos) can modulate TLR7 and TLR8 sensing of site 1 ligands, such as resiquimod (R848), in a sequence-specific manner^29–31^. Counterintuitively, such PS-oligos can have opposing effects on TLR7 and TLR8 sensing, i.e. antagonism and potentiation, respectively, as evidenced in TLR7- and TLR8-expressing HEK 293 cells (HEK TLR7 and HEK TLR8 hereafter) using homopolymers of ≥8 bases of deoxythymidine (dT) (Supplementary Figure 1a)^30,31^.

To establish a detailed mechanistic understanding of how such oligos modulate TLR7/8, we initially tested whether addition of a terminal 5’-_m_U_m_C motif (where _m_ denotes 2’-OMe) could increase TLR8 potentiation of specific 16-20 base (mer) PS-oligos studied previously. This was based on our prior observation that this 5’-end motif was enriched in TLR8 potentiating 2’-OMe PS-oligos and the concept that 2-3 bases were structurally sufficient to interact with site 2 of TLR8 (Figure 1a, Supplementary Figure 1b)^11,29^. While the addition of a terminal 5’-_m_U_m_C motif in oligo #1-UC potentiated TLR8 sensing in both HEK TLR8 and THP-1 monocytic cells, this was not observed in the context of a locked nucleic acid (LNA)-modified oligo (see oligo #2-LNA-UC, Figure 1a, Supplementary Figure 1b). Interestingly, we also observed that mutation of the 5’-end 2’-OMe region of the TLR8-potentiating oligo #660^29^, changing its _m_U_m_C_m_G into a _m_A_m_C_m_G motif in #660-Mut1, ablated TLR8 potentiation (Figure 1a, Supplementary Figure 1b). The _m_U_m_C_m_G motif was directly involved in the TLR8 potentiating activity of oligo #660, as evidenced by the dampening effect of both its partial (oligo #660-Mod) and complete (oligo #660-LNA) LNA modification (Figure 1b, Supplementary Figure 1c). Similarly, 2’-methoxyethyl (MOE) modification of oligo #660’s entire 5’- end region also blunted TLR8 potentiation, indicating a selective activity of 2’-OMe modification on TLR8 potentiation by the _m_U_m_C(_m_G) motif (Figure 1b, Supplementary Figure 1c). A 5-mer oligo reproducing the 5’-end 2’-OMe region of oligo #660 (denoted #660-5) was sufficient to induce significant TLR8 potentiation of R848 sensing, confirming that this was the effector region of this oligo (Figure 1c, Supplementary Figure 1d). Critically, analysis of the three possible 2’-OMe-modified 3-mer oligos encompassing this 5-mer region demonstrated that the _m_U_m_C_m_G oligo was the only sequence to mirror the effect of the 5-mer oligo on TLR8 potentiation (Figure 1d). Potentiation of R848 sensing by _m_U_m_C_m_G, but not _m_C_m_U_m_U, averaged ∼2-fold over a range of R848 concentrations, supporting the concept that it cooperated with site 1 activation (Figure 1e). Three bases was the optimal length for TLR8 potentiation as revealed by the analysis of 2-mer oligos encompassing the same region, which showed that _m_C_m_G also somewhat potentiated TLR8 sensing (Figure 1f).

**Figure 1.**
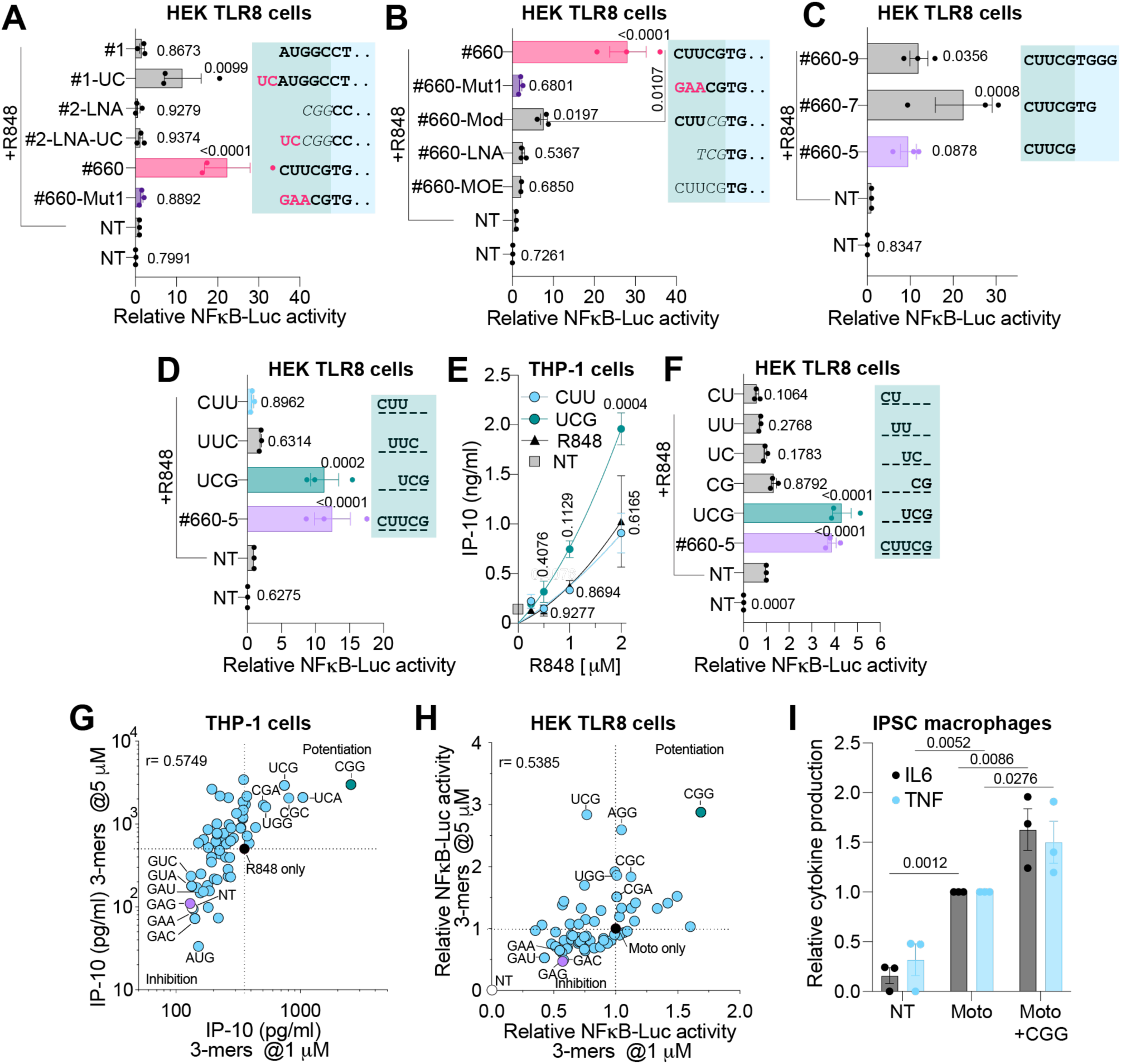
Modulation of TLR8 sensing by 3-mer oligos. (A-D and F) HEK TLR8 cells were pre-treated for ∼30 min with 500 nM (A and B), 2 μM (C), or 5 μM (D, F) of the indicated oligos prior to overnight stimulation with 1 μg/ml of R848 followed by luciferase assay. (A-D and F) Data were background-corrected using the non-treated (NT) condition and are shown as expression relative to R848-only (± s.e.m. and one-way ANOVA with uncorrected Fisher’s LSD tests shown compared to R848-only condition). (B) Unpaired t-test comparing #660 to #660-Mod conditions is shown. (E) Monocytic THP-1 cells were incubated overnight with 1 μM oligo and stimulated with increasing concentrations of R848 (0.250, 0.5, 1, and 2 μg/ml) for 8 h prior to IP-10 ELISA (± s.e.m. and two-way ANOVA with uncorrected Fisher’s LSD tests shown compared to R848-only condition). (G) Monocytic THP-1 cells were incubated overnight with 1 or 5 μM of fully-modified 2’-OMe PS 3-mers and stimulated with 1 μg/ml R848 for 8 h prior to IP-10 ELISA analysis. (H) HEK TLR8 cells were pre-treated for ∼30 min with 1 or 5 μM of fully-modified 2’-OMe PS 3-mers and stimulated with 600 nM of motolimod (Moto) overnight prior to luciferase assay. Data were background-corrected using the non-treated (NT) condition and are shown as expression relative to motolimod only. (I) iPSC-derived macrophages were pre-treated for 30 min with 5 μM of _m_C_m_G_m_G 3-mer prior to stimulation with 400 nM motolimod for 6 h followed by IL-6 and TNF ELISA. Cytokine levels were normalised to the motolimod-only condition (± s.e.m. and two-way ANOVA with uncorrected Fisher’s LSD tests shown compared to the motolimod-only condition). (A-F and I) Data are shown as mean of n=3 independent experiments. (G, H) Data are averaged from 2 or 3 biological replicates for each screen, and the screens at the different oligo concentrations were conducted on independent days. (A-D, F) Green shading represents 5’-end base modification of the oligos as follows: bold pink or black is 2’-OMe, italic is LNA, non-bold is 2’-MOE. DNA bases are shaded in light blue, and ”..” denotes that the sequence is truncated – see Table S5 for full-length sequences.

While these findings aligned with the structural observation that 2-3 RNA bases were sufficient to interact with site 2 of TLR8, the selective potentiating activity of _m_U_m_C_m_G was surprising given the reported inhibitory role of _m_U on TLR7/8 sensing^18,20^. To further define the landscape of TLR8 modulation by short 2’-OMe oligos, we conducted an unbiased screen of all 64 possible 3-mer 2’-OMe PS oligos on TLR8 function in THP-1 and HEK TLR8 cells (Figures 1g, 1h and Supplementary Table S1). Both cell lines revealed a robust TLR8-potentiating activity of 3-mer oligos restricted to a few motifs, including _m_U_m_C_m_G, _m_C_m_G_m_G (the strongest), _m_C_m_G_m_C, _m_C_m_G_m_A, and _m_U_m_G_m_G (Figures 1g, 1h and Supplementary Figure 1e). Potentiation of _m_C_m_G_m_G was independent of the TLR8 agonist used (*e.g.*, R848 and motolimod), and was validated in human induced pluripotent stem cell (iPSC)-derived macrophages, confirming the relevance of its activity in primary-like macrophages (Figure 1i). Unexpectedly, a limited number of 3-mer 2’-OMe-modified oligos robustly inhibited TLR8 sensing, predominantly _m_G_m_A_m_X oligos (X being A, U, G, C) (Figures 1g, 1h), suggesting a complex modulation of TLR8 sensing by short 3-mer oligos resulting in two opposing responses in a motif-dependent manner.

### 2’-OMe-modified 3-mer oligos modulate TLR7 sensing

We have previously reported that 20-mer 2’-OMe-modified oligos generally inhibit TLR7^29^. Importantly, substitution of the three 5’-end 2’-OMe bases in the 20-mer dC-2 oligo with LNA or MOE bases decreased TLR7 inhibition in both HEK TLR7 cells and mouse RAW-264.7 macrophages (Figure 2a and Supplementary Figure 2a). Positing that, similar to TLR8 sensing, the LNA and MOE modifications impeded direct interaction with TLR7, the inhibitory effect of truncated 5’-end fragments of the dC oligo was tested on TLR7 sensing. The results showed that a 5-mer oligo replicating the 5’-end of the dC oligo significantly inhibited human and mouse TLR7 (Figure 2b and Supplementary Figure 2b). This inhibition could be replicated on human and mouse TLR7 with an _m_G_m_U_d_A 3-mer (where _d_ denotes a DNA moiety). It should be noted that 2-mer oligos covering this region were much less inhibitory than _m_G_m_U_d_A (Figure 2c). The fully 2’-OMe 3-mer _m_G_m_U_m_A was also a potent, dose-dependent inhibitor of human TLR7 with significant inhibition still seen with up to 5 times the concentration of R848 used (Supplementary Figures 2c, 2d).

**Figure 2.**
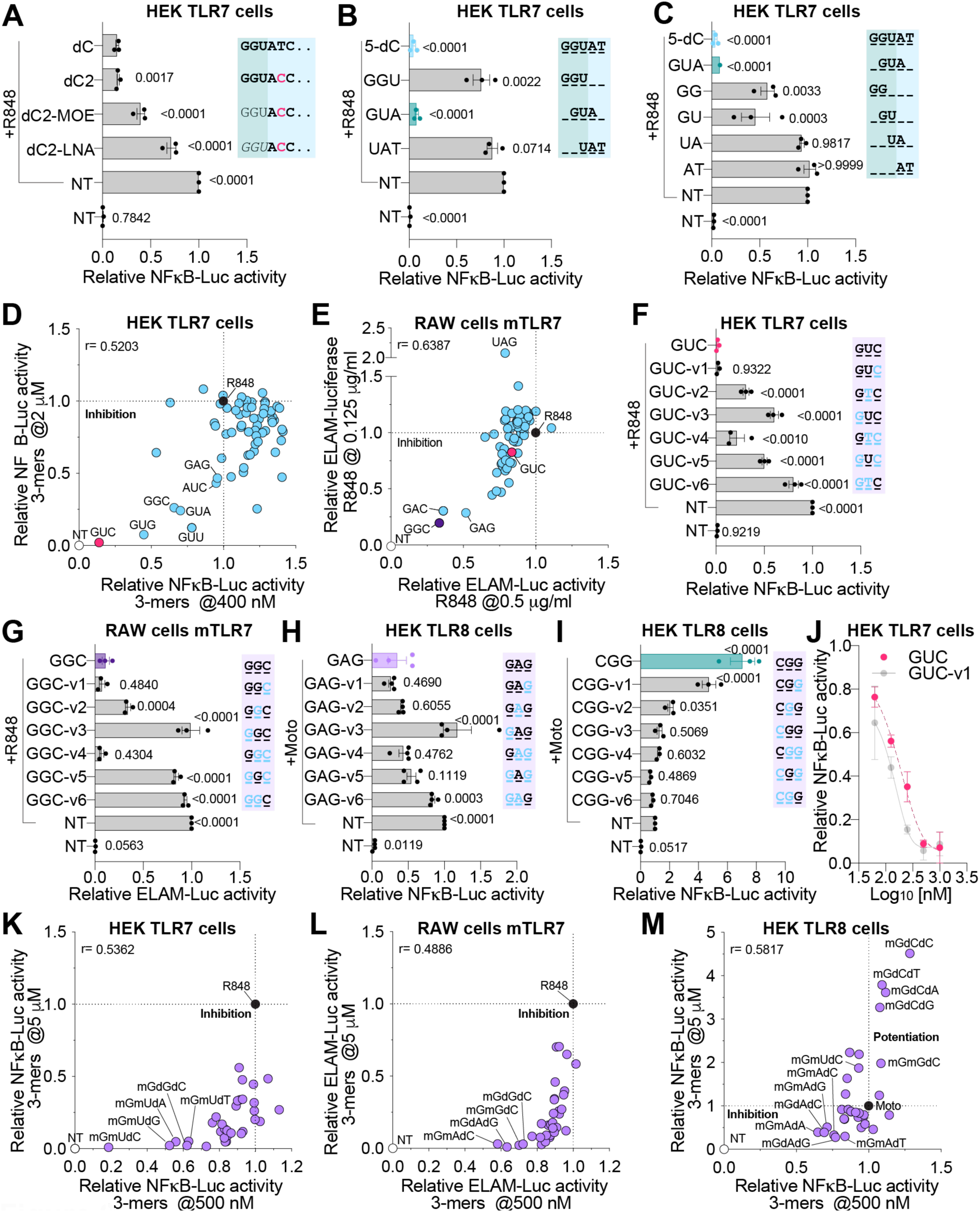
TLR7 inhibition by 3-mer oligos. (A-D, F, J, K) HEK TLR7 cells were pre-treated for ∼30 min with 100 nM (A), 400 nM and 2 μM (D), 500 nM (K), 5 μM (B, C, F, K) or dose-response (0.0625, 0.125, 0.25, 0.5 and 1 μM - J) of the indicated oligos prior to overnight stimulation with 1 μg/ml of R848 followed by luciferase assay. (E, G, L) RAW-ELAM cells were pre-treated with 500 nM (L) or 5 μM (E, G, L) of the indicated oligos prior to overnight stimulation with 0.5 μg/ml (E) or 0.125 μg/ml (E, G, L) of R848 and luciferase assay. (H, I, M) HEK TLR8 cells were pre-treated for ∼30 min with 500 nM (M) or 5 μM (H, Ι, M) of the indicated oligos prior to overnight stimulation with 600 nM motolimod followed by luciferase assay. (A-M) Data were background-corrected using the non-treated (NT) condition and are shown as expression relative to R848/motolimod-only conditions (± s.e.m. and one-way ANOVA with uncorrected Fisher’s LSD tests shown compared to dC+R848 [A], R848-only [B, C], to _m_G_m_U_m_C+ R848 [F], to _m_G_m_G_m_C+R848 [G], to _m_G_m_A_m_G+Moto [H], or to Moto only [I]). (A-C, F-J) Data are mean of n≥3 independent experiments. (D, E, K, L, M) Data are averaged from 3 biological replicates for each screen, and the screens at the different oligo concentrations were conducted on independent days. (A-C) Green shading represents 5’-end base modification of the oligos as follows: bold is 2’-OMe, italic is LNA, non-bold is 2’-MOE. Black or pink DNA bases are shaded in light blue, and ”..” denotes that the sequence is truncated – see Table S5 for full length sequences. (D, E) Fully 2’-OMe-modified 3-mer PS oligos were assessed. (F-I) Black bases in bold denote 2’-OMe modification and light blue bases denote DNA modifications. (K-M) _m_G_m_X_d_X and _m_G_d_X_d_X PS 3-mers were assessed.

These findings prompted us to test the effects of a panel of 64 2’-OMe 3-mers on TLR7 inhibition. The results revealed a strong preferential bias towards _m_G_m_U_m_X 3-mer oligos (X being A, U, G, C) as the most potent inhibitors of human TLR7; _m_G_m_U_m_C being the best, with 75% of the 3-mers showing less than 30% inhibition of TLR7 at the highest dose of 2 μM (Figure 2d). 2’-OMe guanosine was the preferred 5’-end base (13 out of the 16 most inhibitory 3-mers) and, unlike for TLR8, none of the 3-mer oligos robustly potentiated TLR7 sensing (Figures 2d and Supplementary Table S1). Notably, the most potent inhibitor of mouse TLR7 sensing was _m_G_m_G_m_C, followed by _m_G_m_A_m_C and _m_G_m_A_m_G, while the _m_G_m_U_m_C 3-mer oligo inhibited signalling by less than 30%, suggesting structural differences between human and mouse TLR7 underpinning their interaction with the inhibitory 3-mer oligos (Figures 2e).

### 2’-OMe guanosine 3-mer oligos with DNA modifications are potent modulators of TLR7 and TLR8 sensing

The observation that a DNA substitution in the last base of _m_G_m_U_d_A retained inhibitory activity on human TLR7 in Figure 2c, begged the question as to whether the immunomodulatory activity of the 3-mers extended beyond 2’-OMe modification. Interestingly, the TLR7 inhibitory effect was not seen when the 3-mers were fully DNA-modified in screens of 64 such oligos on human and mouse TLR7 (Supplementary Table S1), although minor inhibitory activity of TCT and TTT on human TLR7 sensing was apparent. Similarly, the fully DNA-modified 3-mer oligos did not potentiate TLR8 sensing, although the majority modestly inhibited TLR8 (Supplementary Table S1). To further investigate the importance of each base of the 3-mers in modulating TLR7 and TLR8 function, one or two 2’-OMe bases were systematically replaced with DNA bases (denoted in blue text in the Figure key) in _m_G_m_U_m_C (best human TLR7 inhibitor), _m_G_m_G_m_C (best mouse TLR7 inhibitor), _m_C_m_G_m_G (best human TLR8 potentiator), and _m_G_m_A_m_G (best human TLR8 inhibitor and good mouse TLR7 inhibitor), and we tested their activities on TLR7 and TLR8 sensing (Figures 2f, 2g, 2h, 2i, and Supplementary Figure 2e). For both TLR7 and TLR8 sensing, DNA moieties hampered the activities of 3-mer 2’-OMe oligos when incorporated at the 5’-end, as evidenced by a significant reduction in human and mouse TLR7 inhibition with DNA modification of GUC, GGC, and GAG at this position (see GUC-v3/v6, GGC-v3/v6, and GAG-v3/v6 in Figures 2f, 2g, and Supplementary Figure 2e). Similarly, while both TLR8-potentiating and -inhibitory 3-mers tolerated DNA bases at the third position (see CGG-v1 and GAG-v1) and, to a degree, at the second position (noting a decreased potentiating activity for CGG-v2), DNA modification of their 5’-end base ablated both activities (Figures 2h, 2i - see CGG-v3/v6 and GAG-v3/v6). Interestingly, DNA modification of the 3’-end base instead increased TLR7 inhibitory activity of the GUC-v1 variant over its parental sequence (Figures 2j).

Given that GUC-v4, GGC-v4 and GAG-v4, all of which contain a single 5’-end 2’-OMe guanosine and two DNA bases, retained significant TLR7 or TLR8 inhibitory activity, the activity of a panel of 16 _m_G_m_X_d_X and 16 _m_G_d_X_d_X 3-mer PS-oligos on TLR7 and TLR8 sensing was tested. The analyses of the results revealed close alignment with the initial screens of fully 2’-OMe-modified 3-mer oligos, identifying _m_G_m_U_d_X sequences as the most potent inhibitors of human TLR7, _m_G_m_U_d_C (GUC-v1) being the best (Figure 2k). Similarly, _m_G_m_G_d_C (GGC-v1), _m_G_m_A_d_C (GAC-v1), _m_G_d_G_d_C (GGC-v4) and _m_G_d_A_d_G (GAG-v4) were the most potent inhibitors of mouse TLR7, confirming the tolerance of up to two DNA bases in 3-mer oligos at the 3’-end (Figure 2l). In accordance with the analyses of fully 2’-OMe-modified 3-mer oligos, several DNA modified 3-mers inhibited TLR8 sensing, including _m_G_m_A_d_X (where X is A, T, G or C), and the double-DNA modified _m_G_d_A_d_G (GAG-v4) (Figure 2m and Supplementary Figure 2f). These analyses also revealed novel potentiators of TLR8 sensing with an _m_G_d_C_d_X motif, with _m_G_d_C_d_C (referred to as GCC-v4) being the strongest, outperforming the activity of _m_C_m_G_m_G (Figure 2m and Supplementary Figure 2f). The potentiating activity of GCC-v4 and the inhibiting activity of _m_G_m_A_d_T (referred to as GAT-v1) were validated in iPSC-derived macrophages, where they robustly modulated TLR8 agonist-induced TNF and/or IL-6 production (Supplementary Figures 2g).

### A selective chiral configuration of 2’-OMe 3-mer oligos directly interact with TLR7 and TLR8 to modulate sensing of site 1 agonists

The preceding results demonstrated that a limited number of 5’-end 2’-OMe-modified 3-mer oligos could inhibit TLR7/8 or potentiate TLR8 sensing of their site 1 synthetic agonists, R848 or motolimod, respectively. We also confirmed the capacity of GUC-v1 to inhibit the human TLR7-specific agonist, gardiquimod, and GGC to inhibit gardiquimod, CL075 and ssRNA-driven activation of mouse TLR7 (Supplementary Figure 3a, 3b, 3c). Similarly, GCC-v4 significantly potentiated TLR8 sensing of uridine in iPSC-derived macrophages, while GAG and GAG-v1 significantly inhibited uridine and ssRNA-driven TLR8 sensing in HEK cells (Supplementary Figures 3d, 3e). The activity of the mouse TLR7 inhibitory sequence GGC-v1 was also tested on primary bone marrow-derived macrophages (BMDMs) derived from *Tlr7^Y264H^* gain-of-function mutant mice. This *Tlr7^Y264H^* gain of function mutation results in constitutive engagement of TLR7 through an increased affinity for guanosine^5^. Overnight treatment of *Tlr7^Y264H^* mutant BMDMs with GGC-v1 or the small molecule TLR7/8 inhibitor Enpatoran^32^ led to significant down-regulation of 20 out of the 22 genes regulated by GGC-v1 that were also down-regulated by Enpatoran, as determined by bulk RNA sequencing analyses (Figure 3a, 3b). Importantly, several of the genes confirmed to be significantly down-regulated by both inhibitors with RT-qPCR analyses were previously reported in the top imiquimod-induced genes in a mouse model of psoriatic-like skin inflammation (*e.g.*, Slc13a3, Fpr1, Fpr2, Cd300e)^33^ (Figure 3c).

**Figure 3.**
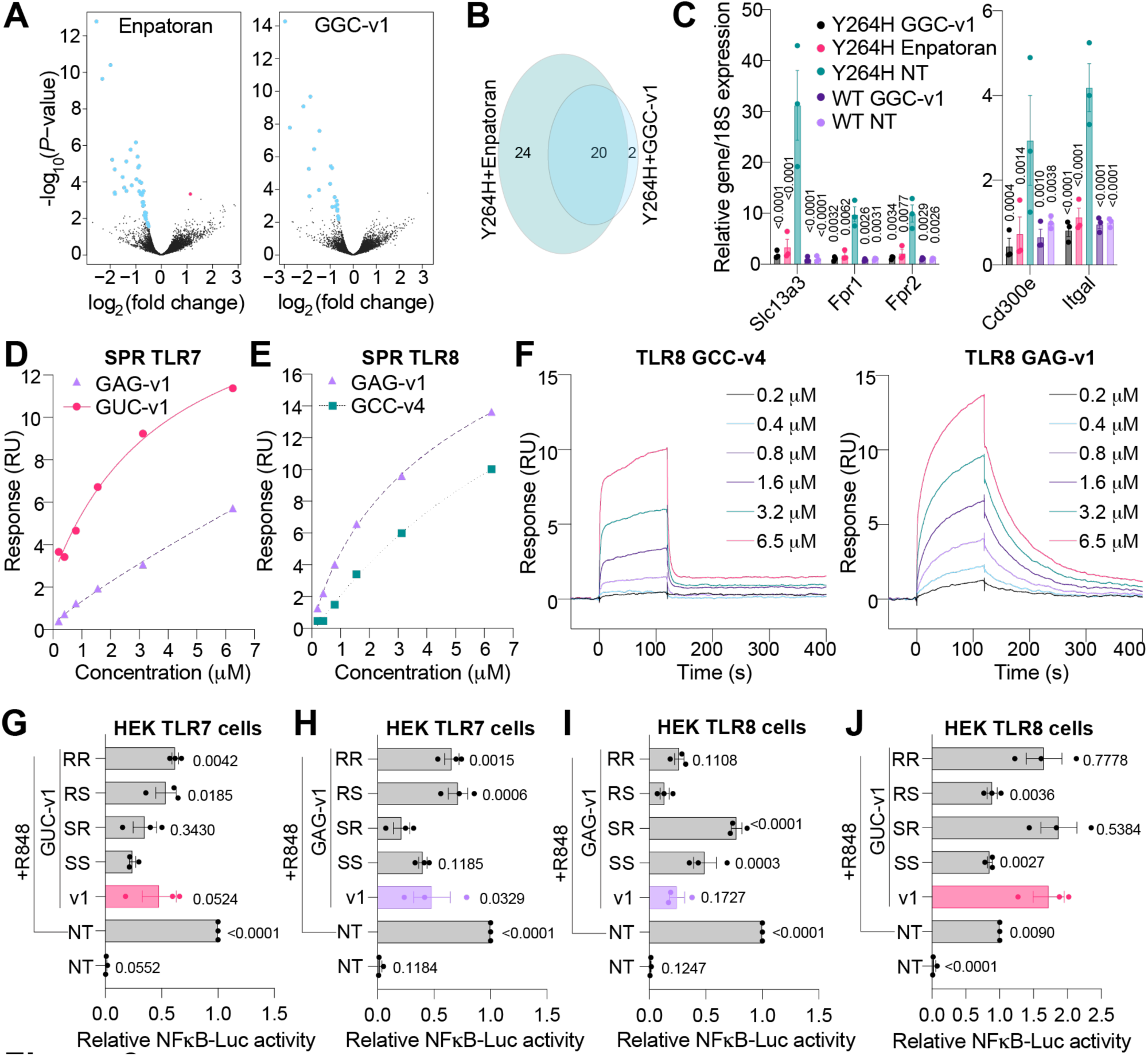
3-mer oligos bind to TLR7/8 to modulate their function. (A-C) Bone-marrow-derived macrophages (BMDMs) from *Tlr7^Y264H^* mice were stimulated for 24 h with 5 μM GGC-v1 or 100 nM Enpatoran prior to RNA purification for RNA-sequencing (A, B) or RT-qPCR analyses (C). (A) Volcano plot of the genes significantly impacted compared to non-treated (NT) condition (blue are down-regulated and red is upregulated) were compared between GGC-v1 and Enpatoran treatments (B). (C) RT-qPCR analyses of *Slc13a3/18S, Fpr1/18S, Fpr2/18S, Cd300e/18S* and *Itgal/18S* in RNA lysates from primary BMDMs from 3 independent *Tlr7^Y264H^* and wild-type (WT) mice. Data are shown relative to the NT condition from WT mice (± s.e.m. and two-way ANOVA with uncorrected Fisher’s LSD tests shown compared to the NT *Tlr7^Y264H^* condition). (D, E, F) Surface plasmon resonance (SPR) analyses of recombinant monkey TLR7 (D) and human TLR8 (E, F) with the indicated concentrations of 3-mers. Data shown are representative of 5-6 independent analyses (Supplementary Table S2). (G-J) HEK TLR7 cells (G, H) or HEK TLR8 cells (I, J) were pre-treated with 200 nM (G) or 5 μM (H, I, J) of _m_G_m_U_d_C or _m_G_m_A_d_G PS 3-mers synthesised as stereopure isomers of RR, RS, SR, or SS configurations, prior to overnight stimulation with 1 μg/ml of R848 followed by luciferase assay. Non-stereopure oligos were included as controls (shown as “v1” conditions). Data were background-corrected using the NT condition and are shown as expression relative to the R848-only condition (± s.e.m. and one-way ANOVA with uncorrected Fisher’s LSD tests shown compared to GUC-v1-SS+R848 [G], to GAG-v1-SR+ R848 [H], to GAG-v1-RS+R848 [I], or to GUC-v1+R848 [J]). (G-J) Data are shown as mean of n=3 independent experiments.

To assess the direct interaction of the lead 3-mer oligos on recombinant human TLR8 and *Macaca mulatta* TLR7, surface plasmon resonance (SPR) was used. The results showed GUC-v1 had an average *K_D_* of 5.6 μM to TLR7, while GAG-v1 (a weaker human TLR7 inhibitor in our cell-based assays), bound to TLR7 with an average *K_D_* of 20.2 μM (Figure 3d, Supplementary 3f and Supplementary Table S2). GUC-v6 and GCC-v4 showed negligible binding to TLR7. Conversely, both GAG-v1 and GCC-v4 bound to TLR8 with good affinities, with averaged *K_D_* values of 4 μM and 8 μM, respectively, while GUC-v6 had negligible binding (Figure 3e, 3f and Supplementary Table S2). Interestingly, the SPR binding profiles of GAG-v1 and GCC-v4 to TLR8 differed substantially – indicative of a different TLR8 binding profile for these oligos, specifically in terms of the kinetics (on- and off-rates) of their interactions with TLR8 (Figure 3f).

These results established the identified lead 3-mer oligos interact directly with TLR7 and TLR8. Importantly, all the oligos tested above were synthesised using phosphorothioate (PS) internucleotide linkages. While reducing nuclease degradation compared to the natural achiral phosphodiester (PO) linkages, the two chiral PS internucleotide linkages in the 3-mers were synthesised in a stereo-random fashion leading to a mixture of four stereoisomers in approximately equivalent quantities. Reasoning that each of these stereochemical configurations might have different binding affinities to TLR7/8, stereopure 3-mer oligos of GUC-v1 and GAG-v1 were synthesized with the four possible configurations (referred to as RR, RS, SR, and SS). The results (Figure 3g, 3h) showed that the RR and RS configurations of GUC-v1 and GAG-v1 displayed significantly less inhibition of human TLR7 in the cell-based assays, compared to the SR and SS variants. Conversely, TLR8 inhibition by GAG-v1 was significantly less with the SR and SS stereoisomers, compared to the RR and RS configurations (Figure 3i). Finally, having observed that GUC-v1 (_m_G_m_U_d_C) acted as a mild potentiator of TLR8 (Figure 2m), its stereoisomers were tested on TLR8 sensing which revealed that the RS and SS configurations blunted potentiation (Figure 3j). Aligning with the SPR data indicating different TLR8 binding profiles, these results supported that TLR8 potentiation and inhibition relate to different configurations of the oligos required for activity (RR and RS for inhibition, and RR or SR for potentiation).

### 2’-OMe 3-mer oligos inhibit TLR7 through interaction with its antagonistic site

Reported crystal structures of *Macaca mulatta* TLR7-RNA complexes indicate the presence of a conserved RNA binding site at the dimerization interface (site 2), where binding of short RNA fragments, including GUCCC, encourages the conformation of the dimer to be in the active form (Figure 4a)^13^. Interestingly, the first three bases of GUCCC RNA are the only ones that directly form interactions with the receptor (Figure 4b, 4c, Supplementary Figure 4a). Given the SPR observations that the 2’-OMe _m_G_m_U_m_C 3-mer oligo binds to TLR7 on its own (Figure 3), we investigated whether this occurred via an increased affinity to site 2 since non-modified 3-mer RNA oligos could not directly bind to TLR7 on their own^12^. Notably, *in silico* CpHMD analyses revealed that, while the truncated 3-mer GUC RNA could stably bind to TLR7 site 2, the 2’-OMe _m_G_m_U_m_C analogue did not remain stably bound to this site. Specifically, the central _m_U base retreated from the conserved TLR7 binding pocket due to a steric clash of the 2’-OMe group with the protein (Figure 4d). This led to a large movement away from the GUC binding site, as seen with the molecule root-mean-square deviation (RMSD) analysis over time and the decreased intermolecular interactions of the 2’-OMe uridine with all the associated TLR7 residues (Figure 4e, Supplementary Figure 4b, 4c). Therefore, the molecular dynamics simulation presented here suggested that the presence of a 2’-OMe uridine group in _m_G_m_U_m_C or GUC-v1 was detrimental to the interaction with TLR7 site 2.

**Figure 4.**
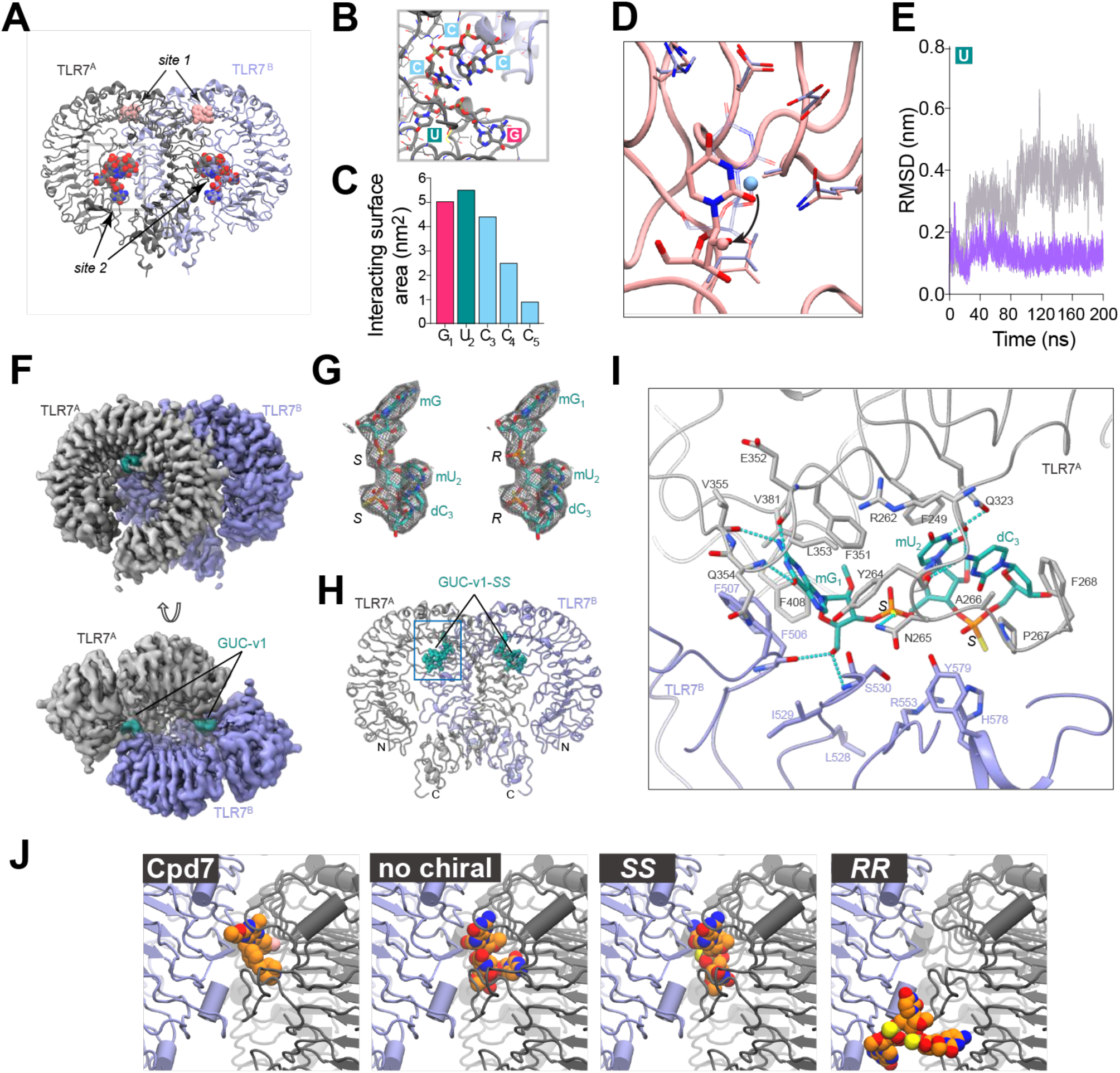
3-mer oligos bind to an antagonistic site in TLR7. (A, B) Crystal structure of TLR7 dimer incomplex with IMQD at site 1 (pink colored ball representation) and GUCCC motif at site 2 (pink)(zoom view in B with details of the RNA binding site shown [PDB: 5ZSE]). The two protomers are colored in dark grey and light blue. (C) Molecular surface area of each nucleotide interacting with protein residues forming the binding site 2, the first three nucleotides contribute the majority of molecular surface (∼80%). (D) The retreat of uracil in _m_G_m_U_m_C from the conserved binding pocket is shown in solid licorice as per MD simulation at pH 5. Conversely, the native uracil from _r_G_r_U_r_C in the binding pocket is shown in transparent licorice. The protein ribbon and sidechains are shown in pink in _m_G_m_U_m_C simulations, and in light blue in _r_G_r_U_r_C simulations. The location of the uracil 2’-residues are indicated by a sphere in each structure. (E) Root mean square deviation (RMSD) of GUC (purple) and _m_G_m_U_m_C (grey) versus time shows that the methylated version moves away from the GUC binding site, indicating the introduced 2’-O-methyl moieties on sugar backbones destabilise the binding of _m_G_m_U_m_C, compared with native GUC. (F) Unsharpened cryo-EM map of the TLR7/GUC-v1 complex shown as surface representations. Densities for the two TLR7 protomers and GUC-v1 are respectively colored gray, purple, and green. (G) Densities around modelled GUC-v1-*SS* (left) and GUC-v1-*RR* (right) shown in mesh representations. The segmented densities were calculated using the surface zone command in ChimeraX. (H) Overall structure of the TLR7/GUC-v1-*SS* complex. Two TLR7 protomers and GUC-v1-*SS* are shown in cartoon and ball-stick representations, respectively. Color schemes are the same as in (F). (I) Close-up view of GUC-v1-*SS* recognition at the antagonistic site. Residues (within 4.5 Å from the ligand) are shown in stick representations and are colored by atom, with the N, O, and S atoms colored by blue, red, and orange, respectively. Dashed lines in cyan indicate hydrogen bonds (cutoff distance < 3.5 Å). (J) The binding pose of TLR7 antagonist, Cpd7, and the best docked poses of _m_G_m_U_m_C with, and without, chiral thiophosphate groups at the antagonistic site.

Positing that the 3-mer oligos could interact with another region of TLR7, cryo-EM analysis of the *Macaca mulatta* TLR7 extracellular domain was performed in the presence of GUC-v1 (racemic mixture of RR, RS, SR, and SS configurations) and the structure of the TLR7/GUC-v1 complex was solved with an overall resolution of 3.0 Å (Figure 4f and Supplementary Table S3). The TLR7/GUC-v1 complex formed a *C*2 symmetric open-form dimer, similar to the previously reported small-molecule antagonist-bound TLR7 structures^34^ (Supplementary Figure 4d). GUC-v1 molecules were clearly observed at the antagonistic site between two TLR7 protomers (Figure 4f). Unlike the closed form of the agonist-bound TLR7 dimer, the two TLR7 protomers in the open form are separated at the C-termini which hinders the proximity of the intracellular TIR domains for activation, thus representing an inhibited state (Supplementary Figure 4d). Although the cryo-EM map may represent the average densities for racemic mixture of GUC-v1 stereoisomers, each stereoisomer could be reasonably fitted to the cryo-EM map (Supplementary Figure 4e). Figure 4g shows the structures of the RR and SS stereoisomers, which are essentially identical in terms of the recognition of the modified nucleotide, with minor variations observed only at the phosphorothioate linkage portion (Figures 4g, 4h, 4i and Supplementary Figure 4f). In support of some isomers binding preferentially to the antagonistic site, separate docking studies conducted without knowledge of the TLR7/GUC-v1 binding site showed that only SS and SR isomers found this site due to improved interactions with S530 (Figure 4j). Hereafter, the representative TLR7/GUC-v1-*SS* complex structure is described because of the relatively stronger inhibitory activity of GUC-v1-*SS* (Figures 3g and 4i).

The 5’-end 2’-OMe guanosine (_m_G_1_) of GUC-v1 deeply inserts into the antagonistic site and makes extensive contacts with TLR7 (Figure 4i). The guanine moiety is stacked by F351^A^ and F507^B^, and is surrounded by bulky aromatic residues including Y264^A^, F408^A^ and F506^B^. The guanine N1 amino, C2 amino, and C6 carbonyl groups form hydrogen bonds with E352^A^ and V355^A^ main chain O atoms, and with the Q354^A^ main chain N atom, respectively. The intimate contacts and hydrogen bonding pattern explains the preference for a guanine base at this position. The 5’-OH group of mG_1_ also forms hydrogen bonds with the F506^B^ backbone carbonyl and S530^B^ backbone amine groups. In addition, the modified 2’-OMe group of mG_1_ points to a small hydrophobic patch formed by F351^A^, V381^A^, and F408^A^ side chains, thereby strengthening the interactions. These structural features are in agreement with the stronger inhibitory effect of the 3-mer oligos with a 2’-OMe guanosine at the 5’-end (Figure 2d). For the phosphate backbone, the first phosphorothioate group forms hydrogen bonds with N265^A^ and S530^B^, and the second phosphorothioate group forms weak electrostatic interactions with R553^B^ and H578^B^. Compared to the stringent recognition of mG_1_, the following _m_U_2_ and _d_C_3_ are loosely recognized. The two pyrimidine rings successively stack onto the F349^A^ side chain and are also surrounded by the P267^A^ and F268^A^ side chains on the opposite side, thereby occupying the entrance of the antagonistic site. Additionally, the N3 amino group of _m_U_2_ and the C4 amino group of _d_C_3_ form hydrogen bonds with the Q323^A^ side chain and with the R262^A^ and Y264^A^ main chain O atoms, respectively. The 2’-OMe group of _m_U_2_ is oriented toward the solvent and positioned between the F349^A^ side chain and the ribose of _d_C_3_.

### 2’-OMe 3-mer oligos modulate TLR7 sensing in vivo

Having established the capacity of _m_G_m_G_m_C and GGC-v1 to block mouse TLR7 signaling driven by ssRNA and chemical agonists, we next investigated the capacity of GGC-v1 to antagonize TLR7 sensing of R848, *in vivo*. Prophylactic intravenous (i.v.) administration of GGC-v1 complexed with the commercial polycationic agent *in vivo*-jetPEI^®^ significantly decreased the splenic induction of several key NF-κB targets driven by intraperitoneal (i.p.) injection of R848 (e.g., *Tnf*, *Il6* and *Il10*), leading to a significant decrease in circulating TNF protein levels in the sera of WT mice (Figures 5a, 5b). Similarly, pre-treatment of the skin of mice with GGC-v1 formulated in 30% F127 Pluronic gel significantly reduced a TLR7-dependent gene signature driven by repeated topical administration of Aldara cream containing imiquimod (including pro-inflammatory *Tnf*, *Cxcl1* and *Il17*, and specific genes reported to be induced in this model e.g. *Fpr1* and *Scl13a3*^33^) (Figure 5c). This reduction in TLR7-driven gene expression in the skin was partially dose-dependent and concurrent with a significant decrease in CD45^+^ immune infiltrates in the skin, and overall decreased skin redness and scaliness (Figure 5d and Supplementary Figures 5a, 5b). Importantly, pre-treatment of the skin with GGC-v1 did not alter the splenomegaly seen in this model, suggesting that its anti-inflammatory effect on TLR7 was localized to the skin (Supplementary Figure 5c, 5d). Collectively, these results established the capacity of GGC-v1 to antagonize TLR7 sensing of its agonists R848 and imiquimod *in vivo*.

**Figure 5.**
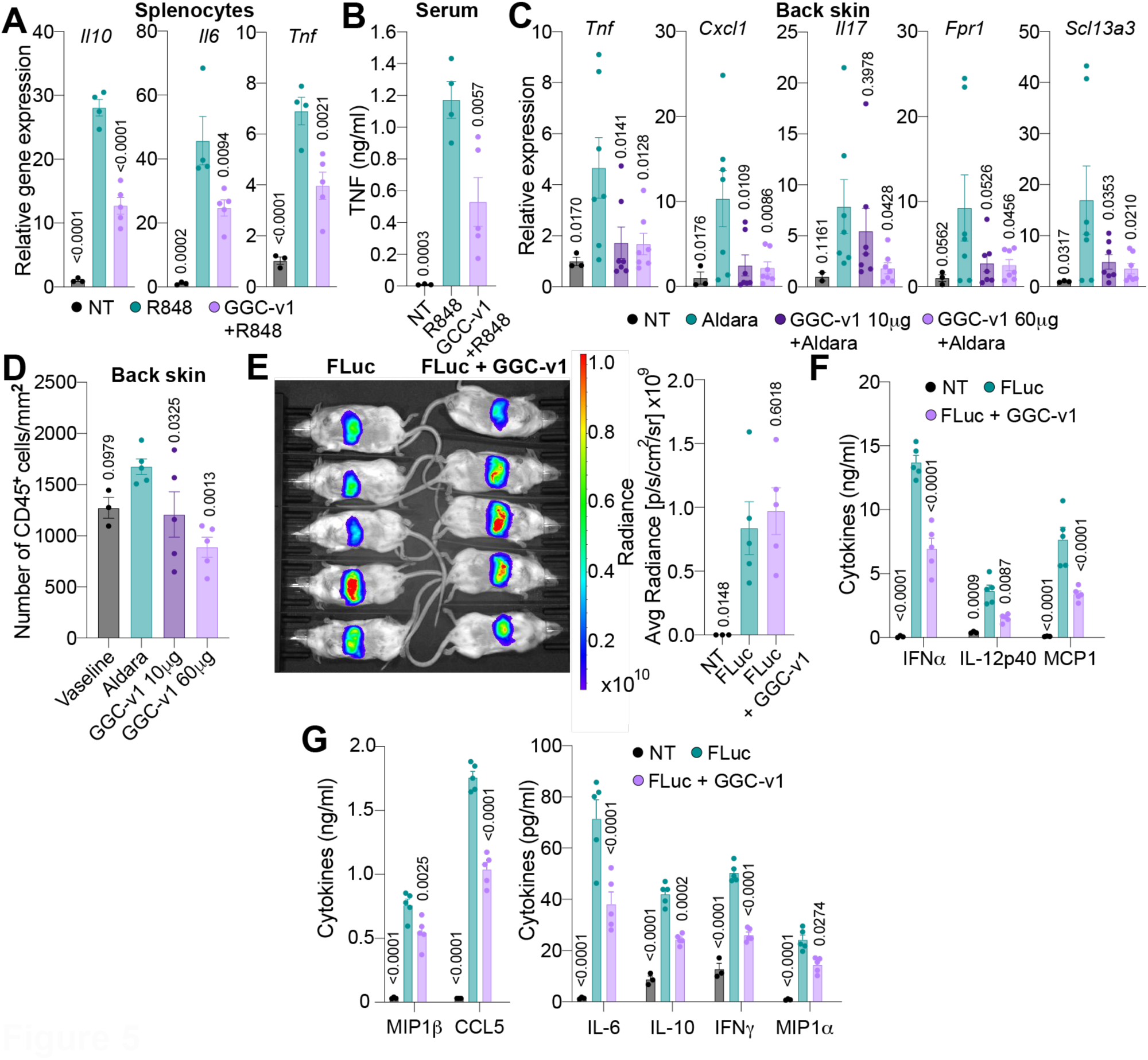
2’-OMe 3-mer oligos antagonise TLR7 function *in vivo*. (A, B) WT C57/BL6 mice were injected i.v. with 200 μg of GGC-v1 conjugated with *in vivo*-jetPEI® for 1 h prior to i.p. injection of 25 μg R848 for 2 h before collection of spleens (A) and sera (B). (A) RT-qPCR analyses of *Tnf/Gapdh*, *Il6/Gapdh* and *Il10/Gapdh* from spleen lysates; data are reported relative to the non-treated (NT) condition. (B) TNF levels were quantified by LegendPlex assay. (A, B) Mean of n=3-5 mice/group is shown (± s.e.m. and one-way ANOVA with uncorrected Fisher’s LSD tests shown compared to R848-only group). (C, D) Aldara cream was applied topically to the back of WT C57/BL6 mice directly following, or not, application of 10 μg or 60 μg GGC-v1 formulated in F127 Pluronic gel for four days. Mice were humanely euthanised and the back skin collected for RNA purification (C) or histology (D). (C) RT-qPCR analyses of indicated genes reported to that of 18S expression, relative to NT mice. Data are representative of 3 independent experiments. (D) CD45^+^ positive cells in the back skin were quantified by fluorescent histology (see Methods). (C, D) Mean of n = 3-7 mice/group is shown (± s.e.m. and one-way ANOVA with uncorrected Fisher’s LSD tests shown compared to Aldara-only group). (E, F, G) WT 129×1/SvJ mice were injected i.v. with LNPs containing 20 μg of FLuc mRNA alone or 17.5 μg FLuc mRNA conjugated to 2.5 μg GGC-v1 (see Methods). (E) IVIS measurement of radiance was conducted at 6 h post-LNP injection and 3-5 min after injection of d-luciferin potassium, and 6 h sera were collected for multiplex cytokine analyses (F, G). Mean of n=3-5 mice/group is shown (± s.e.m. and one-way ANOVA with uncorrected Fisher’s LSD tests shown compared to FLuc-only LNP group).

To determine whether 3-mer oligos inhibited RNA sensing by TLR7 *in vivo*, we studied the activity of GGC-v1 co-administered with 5’-capped T7-synthesised Firefly luciferase mRNA (FLuc) containing unmodified uridine using FDA-approved ALC-0315-based lipid nanoparticles (LNPs) (see Methods)^35^. Following validation that GGC-v1 could be co-packaged with mRNA molecules in LNPs (Supplementary Figure 5e), mice were injected i.v. with LNPs containing Fluc mRNA with or without GGC-v1. While the co-delivery of GGC-v1 did not decrease the expression of FLuc mRNA in the liver, it halved the production of many key pro-inflammatory cytokines in the sera (e.g., IFNα, IFNγ, IL-6, IL-10, IL-12p40, MCP1, MIP1α/β, CCL5 - Figure 5e, 5f, 5g and Supplementary Figures 5f). This is consistent with the concept that the reactogenicity of unmodified mRNAs is at least partially dependent on TLR7^18^ and indicates that the GGC-v1 oligo is capable of dampening TLR7 activation in response to natural ligands *in vivo*. Taken together, these findings demonstrate the capacity of synthetic 2’-OMe 3-mer oligos to functionally modulate TLR7 in biologically relevant animal models.

### RNAs containing select 2’-OMe 3-mer motifs act as natural antagonists of TLR7 and TLR8

Having established the potent inhibitory effect of _m_G_d_G_d_C (GGC-v4) and _m_G_d_A_d_G (GAG-v4) on TLR7 and TLR8, respectively, and given the essential role of _m_G_1_ in the interaction of GUC-v1 with TLR7 (Figure 4i), we reasoned that select _m_G_r_X_r_X motifs naturally occurring in endogenous RNA molecules were likely to modulate TLR7/8 sensing. Focusing on 5’_m_G 3-mers, we screened a panel of 16 _m_G_r_X_r_X-PS oligos on human TLR7 and TLR8 sensing (Figure 6a). Similar to our observations with fully 2’-OMe-modified 3-mers, _m_G_r_U_r_C was one of the most potent human TLR7 inhibitors, while _m_G_r_A_r_G was a strong inhibitor of TLR8 (Figures 6a and Supplementary Table S1). Moreover, additional inhibitors of TLR7/8 were identified across the two receptors, _m_G_r_A_r_A for TLR8 and _m_G_r_G_r_A for TLR7, suggesting that sugar modification of the second base could alter the preferred antagonistic motifs. Notably, the predominant activity of _m_G_r_X_r_X 3-mers was to inhibit TLR7/8 sensing with 8/16 oligos inhibiting both receptors by more than 25%.

**Figure 6.**
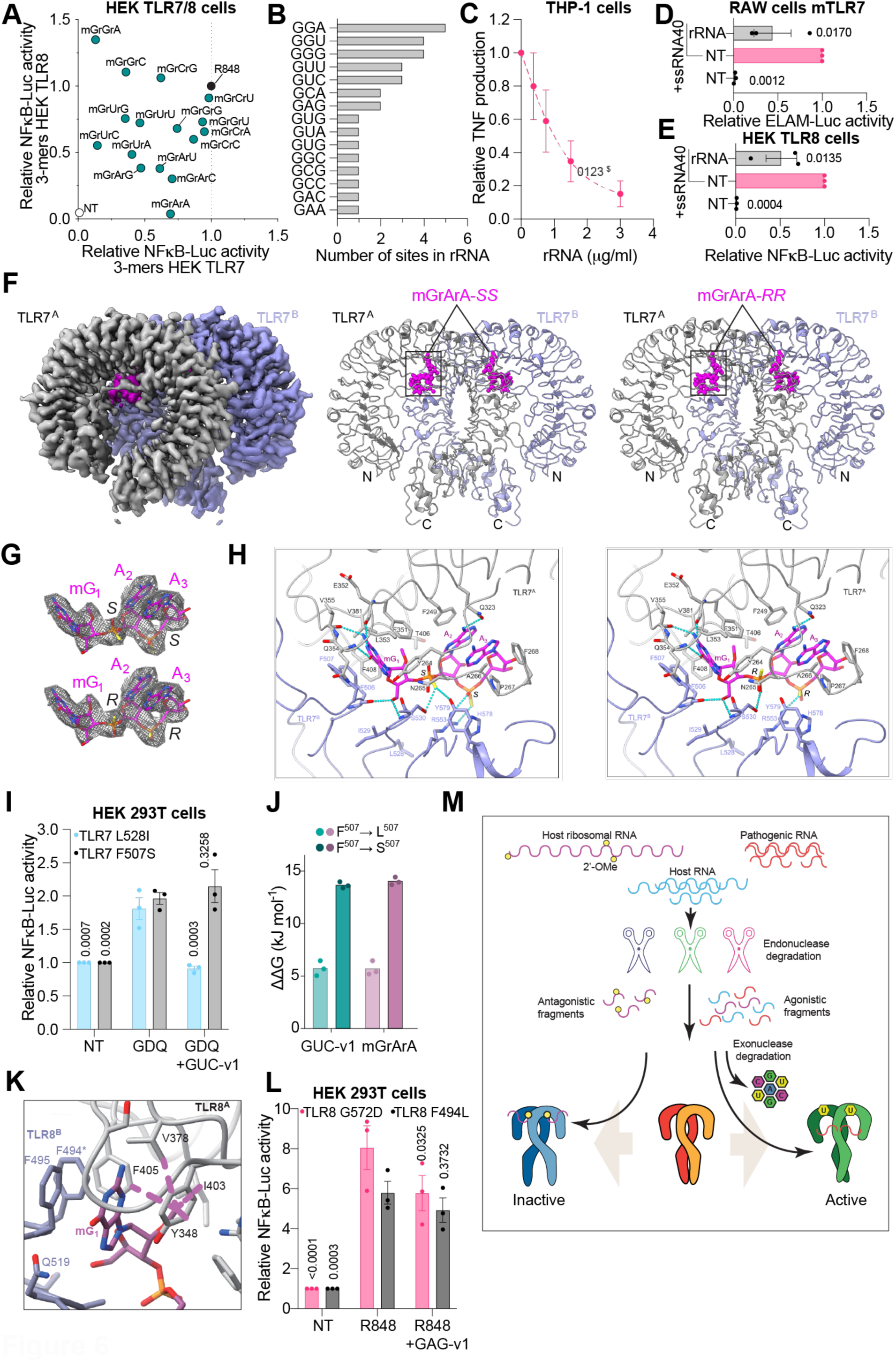
Endogenous 2’-OMe RNA fragments act as natural antagonists of TLR7/8. (A) HEK TLR7 cells (x-axis) and HEK TLR8 cells (y-axis) were pre-treated with 2 μM _m_G_r_X_r_X PS 3-mers prior to overnight stimulation with 1 μg/ml of R848 followed by luciferase assay. Data were background-corrected using the non-treated (NT) condition and are shown as expression relative to the R848-only condition. (B) Cumulative plot of the 2’-OMe G sites previously reported in human rRNA. (C) PMA-differentiated and interferon-γ−primed THP-1 cells were transfected for 6 h with the indicated concentrations of purified rRNA prior to overnight stimulation with 100 nM of transfected ssRNA-40. TNF levels were measured by ELISA and are shown relative to the ssRNA-40-only condition (± s.e.m.). (D) RAW-ELAM macrophages were transfected with 1.5 μg/ml of purified rRNA for 6 h prior to overnight stimulation with 250 nM of transfected ssRNA-40 and luciferase assay. (E) HEK TLR8 cells were transfected with 1.5 μg/ml of purified rRNA for 6 h prior to overnight stimulation with 1 μM of transfected ssRNA-40 followed by luciferase assay. (D, E) Data were background-corrected using the NT condition and are shown as expression relative to the ssRNA-40-only condition (± s.e.m. and one-way ANOVA with uncorrected Fisher’s LSD tests shown compared to the ssRNA-40-only condition). (F) Unsharpened cryo-EM map (left), and structures of the TLR7/_m_G_r_A_r_A-*SS* (middle) and TLR7/_m_G_r_A_r_A-*RR* (right) complexes. The two TLR7 protomers and mGrArA are coloured grey, purple, and magenta, respectively. (G) Densities around mGrArA-*SS* (upper) and mGrArA-*RR* (lower) shown in mesh representations. (H) Close-up view of _m_G_r_A_r_A-*SS* (left) and _m_G_r_A_r_A-*RR* (right) recognition at the antagonistic site. Residues (within 4.5 Å from the ligand) are shown in stick representations and are coloured by atom, with the N, O, and S atoms coloured blue, red, and orange, respectively. Dashed lines in cyan indicate hydrogen bonds (cut-off distance < 3.5 Å). (I) HEK-293T cells co-transfected overnight with the indicated TLR7 mutants, NF-κB-Luc, and UNC93B1 were treated with 1 μM of GUC-v1 for ∼30 min prior to overnight stimulation with 5 μg/ml Gardiquimod (GDQ). (J) The relative binding free energy difference of antagonistic 3-mers (cyan: GUC-v1, purple: _m_G_r_A_r_A) between the wild-type and F507S or F507L mutant (Δ ΔG= Δg_mutant_ - ΔG_WT_). (K) The binding of the best docked pose of PO-_m_G_r_A_r_A to the TLR8 antagonistic site in the inactive dimer (receptor template PDB: 8PFI). Dashed lines indicate the hydrophobic interactions formed among the 2’-O-methyl and the hydrophobic residues highly conserved between TLR7 and TLR8. Carbon atoms and protein ribbons in different protomers are colored in grey and steel blue, carbon atoms in PO-_m_G_r_A_r_A are colored in purple, O, N, and P atoms are in red, blue, and orange respectively. (L) HEK-293T cells co-transfected overnight with indicated TLR8 mutant, NF-κB-Luc, and UNC93B1 were treated with 5 μM of GAG-v1 for ∼30 min prior to 6-8 h stimulation with 1 μg/ml R848. (I, L) Data were background-corrected using the NT condition and are shown as expression relative to the agonist only condition (± s.e.m. and two-way ANOVA with uncorrected Fisher’s LSD tests shown compared to the agonist-only condition). (A) Data are averaged from 3 biological replicates for each screen. (C, D, E, I, L) Data are shown as the mean of n=3 independent experiments. (M) Schematic of natural TLR7 and 8 antagonism. Endosomal nucleic acids from various origins (e.g. host or pathogens) are sequentially processed by endo and exonucleases including RNase T2/2/6 and PLD3/4, respectively. Partial fragments (∼2-3 bases) and single nucleosides (guanosine/uridine) bind to site 2 and site 1 of TLR7/8, respectively. Cooperative binding to site 1 and 2 leads to a closed conformation of the dimers, allowing for downstream signalling. On the other hand, binding of 2-3 base RNA fragments containing 2’-OMe guanosine residues, originating from abundant ribosomal RNA, bind to the antagonistic sites of TLR7 and TLR8 resulting in an inactive open conformation of the dimers. We show that TLR7/8 sensing is kept in check by naturally occurring 2’-OMe modified ribosomal RNA fragments, avoiding autoimmune responses to host RNA in the absence of pathogens.

In mammalian cells, 106 2’-OMe sites have been characterised to-date in 5.8S, 18S, and 28S rRNA^24–26^. Thirty-one of these sites contain a 2’-OMe guanosine, amongst which the most frequent site is the TLR7-inhibitory _m_G_r_G_r_A motif. There are several other frequent inhibitory motifs of TLR7 and TLR8, such as _m_G_r_U_r_C, _m_G_r_A_r_G or _m_G_r_U_r_U (Figure 6b). Critically, purified human rRNA inhibited sensing of the TLR7/8 agonist ssRNA40 in a dose-dependent manner in differentiated THP-1 cells (Figure 6c). rRNA also inhibited ssRNA40 sensing by mouse TLR7 and human TLR8 in RAW cells and in HEK TLR8 cells, respectively (Figures 6d, 6e). Notably, the inhibitory activity of rRNA was also seen on R848-driven TLR8 sensing, confirming that the inhibition operated at the level of the receptor rather than at the level of nuclease processing of the RNA (Supplementary Figure 6a).

Having solved the structure of the TLR7/GUC-v1 complex, and determined that GUC-v1 was a strong antagonist of TLR7, it was of interest to see if the weaker antagonistic _m_G_r_X_r_X 3-mers identified above were interacting similarly with TLR7. Accordingly, we determined the cryo-EM structure of TLR7 in complex with _m_G_r_A_r_A, a more natural 2’-OMe rRNA fragment with modest antagonistic activity, at a resolution of 2.7 Å (Figure 6f and Supplementary Table S3). In this structure, the TLR7/_m_G_r_A_r_A complex also formed a dimer in an open conformation stabilized by _m_G_r_A_r_A binding to the antagonistic site, in a manner similar to the TLR7/GUC-v1 complex (Supplementary Figure 4d). Similar to GUC-v1, _m_G_r_A_r_A also represented racemic mixture of four stereoisomers. We modelled the two representative SS and RR stereoisomers and focused on the TLR7/_m_G_r_A_r_A-*SS* complex (Figure 6g and 6h). The 5’-end _m_G_1_ of _m_G_r_A_r_A-*SS* is recognized by TLR7 in the same manner as the _m_G_1_ of GUC-v1 (Figure 6h, Supplementary Figure 6b), highlighting the strict requirement of 2’-OMe guanosine at the first position. On the other hand, the position of the second and third nucleotide is slightly shifted (Figure 6h, Supplementary Figure 6b), but the phosphorothioate group is similarly recognized: the first phosphate group of _m_G_r_A_r_A-*SS* forms hydrogen bonds with the S530^B^ and Y579^B^ side chains and an electrostatic interaction with R553^B^, and the second phosphorothioate group of _m_G_r_A_r_A-*SS* forms electrostatic interactions with R553^B^ and H578^B^. Two adenine rings of A_2_ and A_3_ are similarly stacked with each other and occupy the entrance of the antagonistic site, as in the TLR7/GUC-v1 complex. The C6 amino group in A_2_ forms a hydrogen bond with the Q323^A^ side chain.

Critically, akin to the interaction with GUC-v1, the guanine moiety is stacked by F507^B^ with the natural _m_G_r_A_r_A-*SS* fragment, indicative of an essential role for F507 in antagonism of TLR7 by natural 2’-OMe rRNA fragments, consistent with reports of SLE patients bearing F507L and F507S gain-of-function (GOF) mutations^4,5^. Accordingly, the antagonistic activity of GUC-v1 on TLR7 was blunted in cells transiently expressing the GOF TLR7 F507S variant, but not the TLR7 L528I mutant (Figure 6i). This is concordant with our *in silico* analyses of the influence of the mutations on the binding free energy of GUC-v1 and _m_G_r_A_r_A obtained using free energy perturbation/Hamiltonian replica exchange molecular dynamics (FEP/H-REMD), indicating that the binding of both is destabilised by mutations at this site (Supplementary Figures 6c-f). As such, the affinity compared to WT of both 3-mers was reduced by more than 200-fold for F507S and 10-fold for F507L (Figure 6j). Noting the conservation of amino acids around residues F506/F507 of TLR7 with TLR8 (aligned to position F494/F495) (Supplementary Figure 6g), and based on prior characterisation of the structure of TLR8 in complex with the small molecule TLR8 antagonist CU-CPT8m^36^, we posited that the F494/F495 residues were also naturally involved in TLR8 antagonism by 2’-OMe rRNA fragments. Although the structure of TLR8 in complex with GAG-v1/ _m_G_r_A_r_A could not be resolved, we successfully docked _m_G_r_A_r_A in the TLR8 antagonistic pocket of an inactive dimer structure^37^, with _m_G_1_ forming direct interactions with F494/F495, and with conserved interactions with the 2’-OMe as seen for TLR7 (Figure 6k and Supplementary Figure 6h), aligning with our prior experimental analyses (Figure 3i). Importantly, we demonstrated that the TLR8 F494L GOF mutation reported in a neutropenic patient^6^ was resistant to GAG-v1 antagonism, unlike another TLR8 GOF mutation G572D (Figure 6l). Collectively, these observations establish that fragments of 2’-OMe-guanosine-modified rRNAs can act as natural TLR7/8 antagonists, and that this natural antagonism is essential for the maintenance of TLR7/8 homeostasis.

## Discussion

A structure of TLR7 bound by a small molecule derived from TLR7 agonists, known as Cpd-7, was recently reported^34^. Notably, unlike agonists of TLR7 that induce a closed form structure, Cpd-7 predominantly induced an open form structure in TLR7, blocking its function^34^. Here, we demonstrate that select 3-mer ssRNA fragments harbouring 2’-OMe guanosine at their 5’-end bind to the same antagonistic site of TLR7^34^, thereby stabilising an open form of the TLR7 dimer and antagonising TLR7 activity. Through extensive functional analyses of all potential variations of _m_X_m_X_m_X, _m_G_m_X_d_X, _m_G_d_X_d_X and _m_G_r_X_r_X 3-base oligos, we established a unique role for 5’-end 2’-OMe guanosine in TLR7 antagonism by select sequences, aligned with the observation that this base led the interaction with TLR7 in the TLR7/GUC-v1 and TLR7/_m_G_r_A_r_A complexes. Our structural analyses revealed that the guanine moiety of our 2’-OMe guanosine oligos is stacked by residues F351^A^ and F507^B^, indicative of an essential role for these residues in the interaction with antagonistic oligos.

Gain-of-function mutations in TLR7 are extremely rare, and have only recently been linked with the onset of systemic autoimmunity, as seen in systemic lupus erythematosus (SLE) patients^3–5^. Given how rare these mutations are, it is notable that SLE patients from two independent families have been reported bearing mutations of F507 in TLR7 (F507S and F507L)^4,5^. While these mutations have been proposed to enhance the agonistic activity of site 1, we demonstrate here that they also compromise the antagonistic activity of GUC-v1 on TLR7 by decreasing the strength of the molecular interaction. This pivotal finding underscores the crucial role of endogenous ssRNA fragments with 5’-guanosine 2’-OMe modification as a checkpoint for TLR7 activation and the potential physiological link between the impaired recognition of endogenous antagonistic RNAs by TLR7 and the onset of systemic autoimmunity.

It is noteworthy that in the structures of TLR7/GUC-v1 and TLR7/_m_G_r_A_r_A complexes presented here, the 3’-end of the oligos faces the solvent region, thereby possibly accommodating longer sequences. This would be consistent with our prior observation that ∼80% of the 192 2’-OMe 20-mer oligos we tested were antagonising TLR7^29^, suggesting that longer sequences can also directly bind to block TLR7 sensing. Analyses of 2’-OMe 2-mer oligos also revealed less TLR7 inhibition compared to 3-mer oligos (Figure 2), indicating that the optimal TLR7 antagonists are >2-base long. These observations collectively suggest that, beyond the case of 3-base RNA fragments structurally evidenced herein, the antagonistic site of TLR7 can be engaged by a compendium of different lengths of 5’-end 2’-OMe guanosine ssRNA degradation products as they are progressively processed by endosomal endo- and exo-nucleases (*e.g.*, RNAse T2, RNAse 2, RNAse 6, PLD3, and PLD4)^15,16,38,39^. In this context it is worth noting that 2’-OMe residues are resistant to RNase T2 and RNase 2 cleavage^16^, suggesting that during the progressive degradation of endosomal ssRNA into single nucleotides, 2∼3-mer fragments containing 2’-OMe residues are likely to be more persistent.

With regards to the origin of natural 2’-OMe-guanosine ssRNA fragments keeping TLR7 in check, we provide evidence that naturally occurring 2’-OMe guanosine RNA motifs present in ribosomal RNA are likely the predominant source. Interestingly, neutrophil extracellular traps (NETs) containing RNAs have been found to engage with TLR8 to exacerbate inflammation, but such NET-associated RNAs contain a limited proportion of rRNA compared to normal cellular RNA (∼25% vs ∼75%, respectively)^40^. This aligns with our proposition that it is the phagocytosed rRNA that modulates TLR7 and TLR8 antagonism. Notwithstanding this, it is likely that other sources of cellular 2’-OMe guanosine RNA are at play in the natural antagonism of TLR7 and TLR8. 5’-Cap-1 mRNA structures and select tRNAs^41^, for example, are 2’-OMe-modified and may also be involved in TLR7/8 antagonism. This is supported by the observation that select bacterial tRNA containing a _m_G_r_G_r_C motif antagonised TLR7^27,28^.

Importantly, the antagonistic mechanism of TLR7 described herein provides a more definitive understanding of TLR7 sensing of RNA degradation products. Structural comparison of TLR7 protomers from the TLR7/_m_G_r_A_r_A-*SS* complex (open inactive form) and the TLR7/GGUUGG complex (closed active form)^13^ showed completely altered Z-loop conformations of TLR7 (Supplementary Figure 6i). As such, _m_G_r_A_r_A-*SS* bound to the antagonistic site directly clashes with the Z-loop region (residues 459-476) in the TLR7/GGUUGG complex, which is essential for shaping the agonistic binding site 2 in its closed configuration. Accordingly, this region is mostly disordered in the TLR7/_m_G_r_A_r_A-*SS* structure. These structural insights demonstrate that ssRNA binding to the antagonistic site and binding to site 2 are mutually exclusive and cannot occur simultaneously. TLR7 sensing therefore results from a competitive activity of uridine-containing ssRNA fragments binding to site 2 synergising with guanosine binding to site 1, working against antagonistic 2’-OMe guanosine RNA binding to the antagonistic site.

In addition to our studies on TLR7, we provide evidence that a similar antagonistic mechanism is at play with TLR8 sensing, also relying on 2’-OMe guanosine RNA fragments. Albeit pending structural validation, our extensive functional analyses of _m_X_m_X_m_X, _m_G_m_X_d_X, _m_G_d_X_d_X, and _m_G_r_X_r_X 3-base oligos indicate a similar observation, with select 2’-OMe guanosine RNA fragments directly antagonising TLR8. The protein region involved in TLR7 antagonism is highly conserved within TLR8, and a rare mosaic mutation of the TLR8 F494 residue (equivalent to TLR7 F506) has been linked to neutropenia in a patient^6^. Notably, the TLR8 F494L gain-of-function protein was refractory to GAG-v1 antagonism in our assays, further supporting a key role for this residue in the antagonistic activity of the lead 2’-OMe guanosine TLR8 antagonists we identified. Given the structural resolution of several small molecule antagonists of TLR8 interacting with F494^32,36,42^, we propose that our 2’-OMe guanosine RNA fragments also bind to this antagonistic site.

However, unlike for TLR7, the regulation of TLR8 sensing by 2’-OMe RNA fragments is not exclusively antagonistic, and we report several sequences, including those with 5’-end 2’-OMe guanosine (*e.g.,* _m_G_r_G_r_A), which instead potentiated TLR8 sensing. Given the different binding profiles of GCC-v4 and GAG-v1 to TLR8 in SPR analyses, we propose that such potentiating 2’-OMe oligos instead bind to site 2, where _r_U_r_G and _r_U_r_U_r_G fragments of RNAs have previously been shown to bind^11^. While further structural studies will be required to confirm this, the identification of very similar 3-base oligonucleotides with opposing activities on TLR8 sensing underlines the complex interplay between agonism and antagonism of this receptor.

There is clear evidence in the existing literature that PS-DNA oligos can inhibit TLR7, an observation we confirmed here with poly(dT) >8-mers^29–31,43,44^. This suggests that, in addition to 3-mer 2’-OMe RNA fragments, DNA fragments may also interact with TLR7/8 antagonistic sites. Our systematic analyses of 3-mer DNA oligos did confirm a modest inhibitory activity of select motifs on TLR8, (*e.g.*, “TTT”), but the inhibition of TLR7 by such 3-mer DNA oligos was very weak overall. Given the capacity of poly(dT) fragments >8-mers to inhibit TLR7, it is therefore likely that the optimal length of DNA antagonists is longer than 3 bases. Through this lens, the endosomal degradation platform generates host DNA and RNA fragments with cumulative antagonistic activities.

In conclusion, our results provide new insight into the mechanisms by which TLR7 and TLR8 activation is normally limited to pathogenic contexts and avoided during homeostatic clearance of apoptotic cells. These findings imply that activation of TLR7 and TLR8 relies on a displacement of natural antagonism (driven by 2’-OMe guanosine), upon accumulation of endosomal agonistic RNA fragments, rather than on the detection of ‘non-self’ RNA features (Figure 6m). The discovery that rare patient mutations in TLR7 and TLR8 residues crucial for this antagonism result in autoimmunity exemplifies the critical role of this natural antagonism in homeostasis. Critically, our findings also evidence the capacity to harness this new anti-inflammatory checkpoint with synthetic 3-mer oligos. Such molecules represent a new class of oligonucleotides with widespread therapeutic applications for the management of inflammatory diseases driven by aberrant TLR7 and TLR8 signalling, including SLE or psoriasis.

## Methods

### Cell culture and reagents

293XL-hTLR7-HA and 293XL-hTLR8-HA stably expressing human TLR7 or TLR8 were purchased from Invivogen and were maintained in Dulbecco’s modified Eagle’s medium plus L-glutamine supplemented with 1× antibiotic/antimycotic (Thermo Fisher Scientific) and 10% heat-inactivated foetal bovine serum (referred to as complete DMEM), with 10 μg/ml Blasticidin (Invivogen). HEK-293T cells^45^ were also maintained in complete DMEM. Human acute myeloid leukemia THP-1 cells were grown in RPMI 1640 plus L-glutamine medium (Life Technologies) complemented with 1x antibiotic/antimycotic and 10% heat inactivated foetal bovine serum (referred to as complete RPMI). THP-1 cells were not differentiated with PMA unless otherwise noted. Overnight THP-1 differentiation was carried out with 20 ng/ml PMA (Merck), and the cells were further primed with 20 ng/ml recombinant IFNγ (Biolegend) for 6 h prior to stimulation with TLR7/8 agonists, as previously reported^46^. RAW264.7-ELAM macrophages^47^ were grown in complete DMEM. All the cells were cultured at 37°C with 5% CO_2_. Cell lines were passaged 2-3 times a week and tested for mycoplasma contamination on routine basis using Mycostrip (Invivogen). Cells were treated with indicated concentration of oligonucleotides or Enpatoran (MedChemExpress) for 20-60 min, prior to R848 (Cayman Chemical), CL075 (Invivogen), Gardiquimod (Invivogen), uridine (Sigma) or Motolimod (MedChemExpress), as indicated. Immunostimulatory ssRNA-40 and B-406-AS-1^48^ ssRNAs, trimer and longer oligonucleotides were synthesised by Integrated DNA Technologies, Syngenis Pty Ltd or Wuxi AppTec, and resuspended in RNase-free TE buffer, pH 8.0 (Thermo Fisher Scientific). For *in vivo* experiments, the oligonucleotides were HPLC-purified and confirmed to be endotoxin free by Limulus Amebocyte Lysate gel-clot method. Sequences and modifications are provided in Supplementary Table S1 and S5. 2’-MOE is moX, 2′-OMe is mX, DNA is dX, RNA is rX, LNA is lX, and phosphorothioate inter-nucleotide linkages are denoted with a *. ssRNAs and purified ribosomal RNA were transfected with DOTAP (Roche). *Tlr7^Y264H^* C57BL/6NCrl mice (used under Australian National University animal ethics, reference A2021/29) have a mutation leading constitutive activation of TLR7 and SLE-like disease^5^. Primary bone marrow derived macrophages (BMDMs) from 9-11 week-old *Tlr7^Y264H^* heterozygous female mice were extracted and differentiated for 5 days in complete DMEM supplemented with L929 conditioned medium as previously reported^49^, prior to 24 h incubation with 5 μM GGC-v1 or 100 nM Enpatoran and total RNA purification for RNA sequencing and RT-qPCRs.

### Human induced pluripotent stem (iPS) cells culture and generation of human iPS-derived macrophages

HipSci HPSI0114i-kolf_2 (ECACC 77650100) iPS line was a gift from Wellcome Trust Sanger Institute and routinely cultured on growth factor-reduced Matrigel (Corning)-coated 6-well plates in mTeSR Plus medium (StemCell Technologies). The iPS line was regularly validated for normal karyotype and negative mycoplasma. Differentiation of human Kolf2 iPS cells toward the macrophage lineage were adapted from ^50^ (H. Hosseini Far et al., *in preparation*).

### Plasmids

pCMV6 vectors expressing Human TLR7 F507S and L528I (gifts from David C. and Crow Y.) and the UNC93B1-mCitrine vector were previously reported^4,51^. pRP[Exp]-mCherry-CMV>{TLR8-F494L} and pRP[Exp]-mCherry-CMV>{TLR8-G572D} expressing GOF TLR8 variants^6^ under the control of a CMV promoter were cloned, amplified and sequence-validated by Sanger-sequencing by VectorBuilder Inc. pCMV6-TLR7-F507S and L528I were purified using an EndoFree® Plasmid Maxi Kit (Qiagen), and were sequence-validated using nanopore whole-plasmid sequencing service from Micromon Genomics Sanger Sequencing Facility (Monash University).

### Luciferase assays

HEK293 cells stably expressing TLR8 or TLR7 were reverse-transfected with pNF-κB-Luc4 reporter (Clontech), with Lipofectamine 2000 (Thermo Fisher Scientific), according to the manufacturer’s protocol. Briefly, 500,000-700,000 cells were reverse-transfected with 200-400 ng of reporter with 1.2 μl of Lipofectamine 2000 per well of a 6-well plate, and incubated for 3-24 h at 37 °C with 5% CO_2_. Following transfection, the cells were collected from the 6-wells and aliquoted into 96-wells, just before oligo and overnight TLR stimulation. Similarly, the RAW264.7 cells stably expressing an ELAM-Luc reporter were treated overnight. As presented in Figure 6, HEK-293T cells were co-transfected with 300 ng or 200 ng TLR7 GOF or TLR8 GOF vectors, respectively, along with 100-150 ng of human UNC93B1-mCitrine^51^ and 50 ng of pNF-κB-Luc4 reporter per well of a 6 well-plate with 1.5 μl lipofectamine 2000. Following overnight incubation, the cells were collected from the 6-wells and aliquoted into 96-wells, just before oligo and overnight (TLR7) or 6-8 h (TLR8) TLR stimulation. In all cases, the cells were lysed in 40 μl (for a 96-well plate) of 1X Glo Lysis buffer (Promega) for 10 min at room temperature. 15 μl of the lysate was then subjected to firefly luciferase assay using 35 μl of Luciferase Assay Reagent (Promega). Luminescence was quantified with a Fluostar OPTIMA (BMG LABTECH) luminometer.

### Cytokine analyses

Production of human IP-10, IL-6 or TNF levels were measured in supernatants from iPSC-macrophages or THP-1 cells using the IP-10 (BD Biosciences, #550926), IL-6 (BD Biosciences, #555220) and TNF (BD Biosciences, #555212) ELISA kits, respectively. Tetramethylbenzidine substrate (Thermo Fisher Scientific) was used for quantification of the cytokines on a Fluostar OPTIMA (BMG LABTECH) plate-reader. All ELISAs were performed according to the manufacturers’ instructions. Concentration of TNF in mouse serum samples (Figure 5b) was quantified using LEGENDplex™ Mouse TNF-α Capture Bead A6 (Biolegend, #740066) as part of the Mouse Anti-Virus Response Mix and Match Panel (Biolgend #740625, #740624, #740623) according to the manufacturer’s instructions. Sample acquisition was performed using a BD LSR-II flow cytometer (BD Biosciences) and the data analysed with the LEGENDplex™ Data Analysis Software Suite (BioLegend).

### RNA and RT-qPCR analyses (Figures 3c and 5c)

Total RNA was purified from *Tlr7^Y264H^* primary BMDMs and using the PureLink RNA Mini Kit (Thermo Fisher Scientific) and DNase-treated using the Purelink DNASE set (Thermo Fisher Scientific). Ribosomal RNA was enriched from 5 μg HEK 293 total RNA obtained with the PureLink kit, using the Ribominus eukaryote kit v2 (Thermo Fisher Scientific) with minor adaptations to the manufacturer’s instructions. Briefly, the beads bound to rRNA were resuspended in 300 μl RNAse free water, and heated 5 min at 70 °C to elute the rRNA. The beads were collected with the DynaMag™ 2 Magnetic Stand (Thermo Fisher Scientific), and the remaining rRNA solution purified further with the PureLink kit.

Random hexamer cDNA was synthesised from isolated RNA using the High-Capacity cDNA Archive kits (Thermo Fisher Scientific) according to the manufacturer’s instructions. RT-qPCR was carried out with the Power SYBR Green Master Mix (Thermo Fisher Scientific) on a QuantStudio 6 Flex RT-PCR system (Thermo Fisher Scientific) with the QuantStudio™ Real-Time PCR Software v1.7.2. Each PCR was performed in technical duplicate and mouse *18S* were used as the reference gene. Each amplicon was gel-purified and used to generate a standard curve for the quantification of gene expression. Melting curves were used in each run to confirm specificity of amplification. Primers used were the following; Mouse 18s: Rn18s-FWD GTAACCCGTTGAACCCCATT; Rn18s-REV CCATCCAATCGGTAGTAGCG; M-F-Slc13a3 GGA AGG CCG ATG CCT CTA TG; M-R-Slc13a3 GGA AGT TGG TGT CGA GGA AGT; M-F-Itgal CCA GAC TTT TGC TAC TGG GAC; M-R-Itgal GCT TGT TCG GCA GTG ATA GAG; M-F-Fpr1 CAT TTG GTT GGT TCA TGT GCA A; M-R-Fpr1 AAT ACA GCG GTC CAG TGC AAT; M-F-Fpr2 GAG CCT GGC TAG GAA GGT G; M-R-Fpr2 TGC TGA AAC CAA TAA GGA ACC TG; M-F-Cd300e TGG GTC TTA CTG GTG CAA GAT; M-R-Cd300e CTT ACA CTG ACC GAT GGA TCA C; Ms Cxcl1-F1 CCT TGA CCC TGA AGC TCC CT; MsCxcl1-R1 CAG GTG CCA TCA GAG CAG TCT; mIL17aF ACC GCA ATG AAG ACC CTG AT; mIL17aR TCC CTC CGC ATT GAC ACA; MsTnfaF1 CAA AAT TCG AGT GAC AAG CCT G; MsTnfaR1 GAG ATC CAT GCC GTT GGC.

### RNA sequencing

Libraries were generated using an in-house multiplex RNA-seq method (version 01/09/2021; Hudson Genomics Facility) adapted from^52^. Briefly 25 ng of total RNA from each sample was tagged with an 8 bp sample index and 10 bp unique molecular identifier (UMI) during initial poly(A) priming with the addition of a template-switching oligo. After cDNA amplification Illumina P5 adaptors with distinct i5 indexes were added by tagmentation by Nextera transposase and PCR. The final libraries were pooled based upon size-adjusted qPCR and single-end sequencing was performed on a NextSeq 2000 run using a P3 50 cycle kit (cDNA reads generated 61 nt). Base calling was performed using Dragen BCLConvert (v3.7.4).

### RNA-seq analysis

RNA-seq analysis was performed in R (v4.1.0)^53^. The scPipe package (v1.14.0)^54^ was employed to process and de-multiplex the data. Read alignment was performed on R1 FASTQ files using the Rsubread package (v2.6.1)^55^. An index was built using the Ensembl *Mus musculus* GRCm39 primary assembly genome file and alignment was performed with default settings. Aligned reads were mapped to exons using the sc_exon_mapping function with the Ensembl *Mus musculus* GRCm39 v104 GFF3 genome annotation file. The resulting BAM file was de-multiplexed and reads mapping to exons were associated with each individual sample using the sc_demultiplex function, taking the UMI into account, and an overall count for each gene for each sample was generated using the sc_gene_counting function (with UMI_cor = 1). Additional gene annotation was obtained using the biomaRt package (v2.48.3)^56^ and a DGEList object was created with the counts and gene annotation using the edgeR package (v3.34.0)^57^. A design matrix was constructed incorporating the treatment group and donor mouse (BMDMs from n=3 mice were used). Lowly expressed genes were removed using the filterByExpr function and normalisation factors were calculated using the TMM method^57^. Counts were transformed for differential gene expression analysis using the voom method^58^ and a linear model was fit using the edgeR voomLmFit function. Comparisons were made for each treatment group compared to non-treated controls using the contrasts.fit function and empirical Bayes moderated t-statistics were calculated using the eBayes function^59^. Differentially expressed genes were determined using a false discovery rate (FDR) adjusted p value < 0.05. Volcano plots were made using the log2 fold changes of all genes versus the -log10 p value across all samples as calculated during this step.

The original multiplexed R1 FASTQ files were deposited along with the UMI counts in the NCBI Gene Expression Omnibus (GEO) with accession GSEXXXXX.

### Recombinant TLR7 and TLR8 proteins

Recombinant TLR7 and TLR8 extracellular domains were prepared as previously described^10,12^. For TLR8, *Homo sapiens* TLR8 extracellular domain fused to a C-terminal thrombin-cleavage sequence followed by a protein A tag was constructed into the pMT-BiP-V5-His vector (Thermo Fisher Scientific). For TLR7, *Macaca mulatta* TLR7 extracellular domain (N167Q, N399Q, N488Q, N799Q mutations, and residues 440-445 (SEVGFC) replaced by a thrombin-cleavage sequence (LVPRGS)) was constructed into the pMT-BiP-V5-His vector. For expression, stably transfected *Drosophila* S2 cells (Thermo Fisher Scientific) were cultured in EXPRESS FIVE SFM medium (Thermo Fisher Scientific). Protein expression was induced by addition of 0.5 mM copper(II) sulfate. Secreted proteins were purified using IgG Sepharose 6 Fast Flow affinity resin (Cytiva) and further subjected to thrombin cleavage and gel-filtration chromatography. Purified proteins were used for SPR and cryo-EM experiments.

### Surface plasmon resonance (SPR)

Using a Biacore T200 (Cytiva), TLR7 and TLR8 were immobilised onto a Series S sensor Chip CM5 (GE) by amine coupling to a level of ∼7800 RU and ∼6100 RU respectively as per the manufacturer’s instructions. Briefly, at 25°C, with a flowrate of 10 µl/min, flow cells were activated with injection of 0.2 M EDC + 0.05 M NHS for 420 s. TLR7 (50 µg/ml in 10 mM acetate pH 4) was coupled to flow cell 2 (450 s, 2 ul/min), and TLR8 (100 ug/ml in 10 mM Acetate pH 4) was coupled to flow cell 3 (500 s, 2 ul/min). Unreacted NHS was blocked on all flow cells with an injection of 1 M ethanolamine-HCl pH 8.5 for 420 s. Flow cell 1 was used as the reference, activated and blocked as described above. Immobilisation buffer was 20 mM HEPES, 150 mM NaCl, pH 7.5. The SPR run was performed at 20 °C with 10 mM Mes, 150 mM NaCl, pH 5.5 as the running buffer. All oligonucleotides were solubilised in running buffer, and concentration confirmed via the calculated extinction coefficient at A260 nm. A 6-point dilution series between 0.2 – 6.25 uM was performed for all oligonucleotides. Samples were injected for 120 s with a dissociation time of 300 s at a flow rate of 40 µl/min. Surface regeneration was performed by 2 x 60 s injections of 2 M NaCl at 30 µl/min. Data were analysed using Biacore T200 Evaluation Software. Affinity constants were determined by fitting the data using affinity analysis with a 1:1 binding model. For TLR7:GAG-v1 and TLR8:GCC-v4 interactions, the data were fitted with Rmax fixed based on the TLR7:GUC-v1 and TLR8:GAG-v1 datasets respectively.

### Cryo-EM analysis, data processing and model building

Purified recombinant TLR7 protein (0.2 mg/ml) was mixed with GUC-v1 (mixture of RR, RS, SR, and SS configurations, 0.17 mg/ml) (WuXi AppTec) or _m_G_r_A_r_A (0.3 mg/ml) (WuXi AppTec) using a dilution buffer containing 20 mM sodium citrate tribasic dihydrate, pH 5.0 and 150 mM NaCl. For each sample, a 3-μl aliquot was applied onto a glow-discharged QUANTIFOIL® R 1.2/1.3 on Cu 300 mesh grid + 2 nm C Holey Carbon Films with ∼ 2 nm continuous carbon on top (QUANTIFOIL, #C2-C14nCu30-50). The grids were blotted for 2.0 s in 100% humidity at 6°C and plunged into liquid ethane using a VitrobotMkIV (Thermo Fisher Scientific). Cryo-EM movies were recorded by using a Titan Krios G4 microscope (Thermo Fisher Scientific) running at 300 kV and equipped with a Gatan Quantum-LS Energy Filter (GIF) and a Gatan K3 camera in the electron counting mode at the Cryo-EM facility in the University of Tokyo (Tokyo, Japan). Imaging was performed at a nominal magnification of 105,000×, corresponding to a calibrated pixel size of 0.83 Å/pixel. Each movie was recorded for 1.6 s and subdivide into 48 frames with an accumulated exposure of about 48 e^−^/Å^2^. EPU software (Thermo Fisher Scientific) operated at the fast acquisition mode was used for data collection.

For processing the TLR7/GUC-v1 dataset in RELION v4.0.1^60,61^, 6615 raw movie stacks were motion-corrected using MotionCor2 in RELION’s own implementation^62^. The CTF parameters were determined using the CTFFIND4 program^63^. Particles were picked using the auto-picking program. Multiple rounds of 2D classification, 3D classification, 3D auto-refine, Post-processing, CTF refinement and Bayesian polishing were performed to select particle stacks (173,248 particles) for the final 3D auto-refine, which yielded an overall resolution of 3.0 Å. For processing the TLR7/mGrArA dataset in cryoSPARC v4.0.3^64^, 3111 raw movie stacks were motion-corrected using the patch motion correction, and the CTF parameters were determined using the patch CTF estimation. Particles were picked using the blob picker. Multiple rounds of 2D classification, ab initio reconstruction and heterogeneous refinement were performed to select particle stacks (1,446,780 particles) for final 3D refinement using non-uniform (NU) refinement^65^, which yielded an overall resolution of 2.7 Å. To obtain sharpened 3D maps, *B*-factor sharpening was applied to the 3D maps from the consensus refinement. More detailed cryo-EM image processing is provided in Supplementary Figure 7.

For model building, the TLR7/Cpd-7 structure (PDB 6LW1) was used as the initial model by fitting into the cryo-EM maps using the fit in map program in Chimera software^66^. Ligand restraints (CIF files) were generated using the JLigand software. The iterative cycles of manual model building in COOT^67^ and real-space refinement in Phenix software^68,69^ were continued until the structures converged with reasonable geometric parameters and map-to-model fit. Both the unsharpened and the *B*-factor sharpened 3D maps were referred during refinement. The unsharpened 3D maps were selected as the main maps and have been deposited in the Electron Microscopy Data Bank. The atomic coordinates have been deposited in the Protein Data Bank. Statistics for data collection and structural refinement are summarized in Supplementary Table S3. Structure representations were generated using ChimeraX^70^.

### Molecular dynamics simulations docking studies

The 2.2 Å crystal structure of monkey TLR7 in complex with IMDQ and GGUCCC (PDB ID: 5ZSE)^13^ was used to create the homology model of wild-type human TLR7 active dimer complex using MODELLER (version 10.4)^71^, and the cryo-EM structures of GUCv-1 and _m_G_r_A_r_A bound complex in this study were used as the templates to create the corresponding inactive human TLR7 models. The locations of crystal water molecules were refined and added using COOT^67^ and phenix^72^. Only the nucleotides GUCCC or GUC with clear electron density were modeled in the active complex. The N-glycosylation sites on known asparagine residues were determined using the most common types on each site after checking all available electron density maps. All simulation systems were prepared using the CHARMM-GUI Solution Builder server ^73^, with N- and C-terminal residues patched as acetylated N terminus and methylamidated C terminus, respectively. The 2’-OMe modifications on RNA sugar groups were added and structurally refined using IQmol. For non-free-energy-perturbation simulations, the constant pH molecular dynamics (CpHMD) were conducted at pH 5; the simulation cells comprised approximately 320,000 atoms, of which approximately 100,000 were water molecules, in a box of dimensions 15 × 15 × 15 nm^3^. TIP3P water parameters were used to solvate all systems. The environmental pH value in CphMD was set at value of 5.0, and 450 buffer particles were added to tackle the net charge changes due to the possible transition of protonation states of titratable residues (aspartate, glutamate and histidine). Sufficient Na^+^ and Cl^−^ ions were introduced by replacement of water molecules to bring the systems to an electrically neutral state at an ionic strength of 0.15 M. Specific parameters for describing the interactions involving titratable residues or buffer particles for CpHMD were taken from CHARMM36* all atom force field ^74,75^. For free-energy-perturbation calculations, the initial protein structures and hybrid topology files were obtained using pmx toolkit^76^, where only the residue F507 in the wild-type structure was replaced by hybrid residue F2S or F2L after being superimposed with the pmx made structure lacking these chemical modifications. The protein topology files and force field parameter file generated by CHARMM-GUI using charmm36m all atom force field^77^ were modified according to the pmx hybrid topology files to include the engineered residue F2S or F2L and define possible interactions.

All simulations were performed using the GPU-accelerated GROMACS software package (version 2023.1 and version 2021 with CpHMD module)^78^. CGenff program was used to parameterize the small ligand^79^. For CphMD, steepest descent energy minimization was conducted followed by two sequential steps of equilibration (500 ps in NVT ensemble and 500 ps in NPT ensemble) with a gradual decrease in the restraining force applied to protein atoms and ligands. The LINCS algorithm^80^ was applied for resetting constraints on covalent bonds to hydrogen atoms, which allowed 2 fs time steps for MD integration during the entire simulation. The particle-mesh Ewald algorithm^81^ was used for calculating electrostatic interactions within a cut-off of 12 Å, with the Verlet grid cut-off-scheme^82^ applied for neighbour searching, using an update frequency of 20 and a cut-off distance of 12 Å for short-range neighbours. A 12 Å cut-off was applied to account for van der Waals interactions, using a smooth switching function starting at 1.0 nm. Periodic boundary conditions were utilized in all directions. During the equilibration stages, the temperature was maintained at 303.15 K using a Berendsen-thermostat^83^ with a time constant of 1.0 ps. Protein with ligands and ion-water groups including buffer particles were treated independently to increase accuracy. The pressure was maintained at 1.0 bar by isotropic application of a Berendsen-barostat^83^, with a time constant of 5.0 ps. During production of molecular dynamics, the temperature was maintained at 303.15 K using a v-rescale-thermostat^84^ with a time constant of 1.0 ps, and the pressure was maintained at 1.0 bar using the Parrinello–Rahman-barostat^85^ isotropically, with a time constant of 5.0 ps and compressibility of 4.5 × 10^−5^ bar^−1^. The active human TLR7 dimer bound with GUCCC, GUC and _m_G_m_U_m_C at site 2, plus inactive human TLR7 dimer bound with GUC-v1 and its F507S mutant bound with PO-GUC-v1 were studied by running for 200 ns in 5 replicates, respectively, and a total of 5 µs long simulations were conducted.

For FEP, the Hamiltonian replica-exchange molecular dynamics (H-REMD) was conducted to improve the convergence of free energy calculations. Free energy module in GROMACS was turned on using parameter lambda to interpolate between topology A (lambda=0) to topology B (lambda=1). Soft-core potentials are used with an alpha parameter of 0.5 and sigma parameter set by 0.3 nm, and the power for lambda in the soft-core function was 1. A total of three independent steps were applied to connect the topology A (Phe) and topology B (Ser or Leu). In the first set only the partial charges on residue Phe507 were tuned by having six discrete replicas in which the lambda was 0.0, 0.2, 0.4, 0.6, 0.8 and 1.0, respectively, the residue type was still phenylalanine, and all partial charge values were zero while lambda_step1_=1. In the second set, the partial charge on each atom including dummy atoms was kept zero while the vdw, mass, bonded and restraint terms of those atoms were tuned by having 16 discrete replicas in which the lambda was 0.000, 0.067, 0.134, 0.201, 0.268, 0.335, 0.402, 0.469, 0.536, 0.603, 0.670, 0.737, 0.804, 0.871, 0.938 and 1.000, respectively, and the residue could be recognized as a serine whereas all partial charge values were zero while lambda_step2_=1. In the third set, only the partial charge on each atom was tuned again by having six discrete replicas in which the lambda was 0.0, 0.2, 0.4, 0.6, 0.8 and 1.0 to bring the proper partial charge back to the serine residue, an intact serine was define while lambda_step3_=1. All lambda points were written out by having the calc-lambda-neighbors value of -1. Stochastic dynamics integrator was used during FEP. The attempt replica exchange periodically with every 1000 steps of calculation. Both apo form of inactive dimer and antagonistic RNA 3-mer (PO-GUC-v1 or PO-_m_G_r_A_r_A) binding dimer were used to close the thermodynamics cycle to obtain the relative binding free energy difference. Each set of H-REMD was repeated three times in which each replica was run for 50 ns, in total of 25.2 µs in FEP. Convergence of the results was assessed by plotting the free energy as a function of simulation time. The protonated states of all aspartate, glutamate and histidine residues were pre-assigned using the results from CpHMD residue titration calculations, and using a range of pH from 2.0 to 10.5 with an interval of 0.5 (Supplementary Table S4). The system was simulated at each environmental pH for 50 ns in 5 replicates, resulting in overall simulation length of 4.5 µs.

### Molecular docking

Flexible docking was conducted to investigate the potential of inhibitory RNA 3-mers to bind at the antagonist binding site in the inactive TLR7 (PDB ID: 6LW1) and TLR8 (PDB ID: 8PFI). Autodocktools^86^ was used to parameterize the RNA ligands (PO-mGmUmC and PO-mGrArA) and protein flexible and rigid components. Autodock VINA^87,88^ was used to conduct the flexible docking with exhaustiveness of 32, the grid box has a dimension of 50 Å × 50 Å × 50 Å centering at the geometry center of known antagonist (removed before docking in progress). The flexible residues chosen in TLR7 docking experiment were R262^A^, Q323^A^, F349^A^, P351^A^, R378^A^, V381^A^, F408^A^, F506^B^, F507^B^, and flexible residues chosen in TLR8 docking experiment were F261^A^, Y348^A^, V378^A^, I403^A^, F405^A^, E427^A^, F494^B^, F495^B^, R541^B^. The inhibitory RNA binding poses with mG_1_ docked at the binding pocket (if captured) were always the pose with best calculated affinity.

### Co-encapsulation of GGC-v1 with FLuc mRNA in LNPs

1 mg of CleanCap® FLuc mRNA (TriLink; L-7602) was pre-mixed by pipetting, or not, with 0.2 mg GGC-v1 (5:1 ratio by weight) before encapsulation in lipid nanoparticles (LNPs) with the composition of III-3: IVa PEG-lipid: DSPC: Cholesterol = 47.4:10:40.9:1.7 (by molar ratio) using a Nanoassemblr Ignite (PrecisionNanoSystems). Resulting LNP formulations were dialyzed against 20 mM pH 7.4 Tris buffer and diluted with PBS after dialysis. Concentrations of the LNP-formulated mRNA samples were adjusted to 0.2 g/L. LNP particle size, polydispersity index (PDI), zeta potential, and mRNA encapsulation rate (Ribogreen assay) were assessed and comparable for both formulations generated. LC-MS/MS and RT-qPCR were used to quantitate GGC-v1 and FLuc mRNA, respectively, in the LNPs. FLuc primers used were: FLuc Forward primer: 5’-CCACATCGAGGTGGACATCA-3’; FLuc Reverse primer: 5’-AGCACGGGCATGAAGAACTG-3’. The RT-qPCR analyses were performed on a QuantStudio® 5 Real-Time PCR System (Applied Biosystems) and data analysed using QuantStudio Design and Analysis Software v1.5.2.; LC-MS/MS analyses were conducted on Waters Acquity premier UPLC system (Waters Corporation, Milford, MA) coupled QTRAP 6500 mass spectrometer (Sciex, Framingham, MA) and analysed with Analyst Software v 1.6.3.

### Mouse studies

All the animal experiments complied with relevant local ethical regulations.

### Aldara-driven skin inflammation model

These experiments were approved in advance by an Animal Ethics Committee at Monash Medical Centre (MMCB/2022/18 and MMCB/2023/19) and were carried out in accordance with “Australian Code of Practice for the Care and Use of Animals for Scientific Purposes. Eight weeks old C57Bl/6J female mice were used in these experiments and were housed in SPF (Specific Pathogen Free) conditions. Mice were anaesthetized for 1-2 minutes with a vaporiser machine (oxygen flow rate of 1-4 litres/min) with fresh oxygen and 5% isoflurane. Upon induction of anesthesia, back hair removal was done using sensitive hair removal cream (Nair). A small drop of Nair cream was applied to mouse’s upper back using a cotton bud and smeared to cover a 2cm^2^ patch. The Nair cream was allowed to sit for ∼one minute before being wiped off in a nose-tail motion using a sterile cotton bud to remove hair from the back skin. Thereafter, mice were treated topically to the hairless back skin with 10 μl of 10-60 μg of highly pure GGC-v1 oligonucleotide (>99.4%) resuspended in sterile PBS, combined with 20 μl of ice cold 30% Pluronic F-127 gel solution in sterile PBS (Sigma), and allowed to absorb for 1 min. 50 mg of Aldara cream (containing imiquimod, 5% w/v) was subsequently applied to the back skin using a cotton bud. One cohort of mice was treated with Vaseline cream as vehicle control for the Aldara cream. These treatments were repeated daily for 4 consecutive days. Mice were scored daily for back erythema (redness) and skin scaling, as previously reported^89^. Spleens and skin samples were collected at the end of the experiment (day 5).

### Histology and immunofluorescence

Immunofluorescence staining was performed using the Opal staining kit (Akoya Biosciences) as described previously^90^. Skin tissue sections were deparaffinised, and antigen retrieval was performed by microwave treatment. Endogenous peroxidases were quenched by treating the tissue sections with 3% H_2_O_2_ for 10 min. Tissue sections were incubated in blocking buffer (Akoya Biosciences) for 10 min at room temperature. After this, sections were incubated overnight with CD45 (D3F8Q) antibody (1:200 dilution in SignalStain Antibody Diluent, Cell signalling Technology) at 4°C and incubated with secondary HRP (SignalStain Boost IHC Detection Reagent, Cell signalling) for 30 min at room temperature. The sections were incubated with Opal 570 working solution (Akoya Biosciences) for 10 min at room temperature and counterstained with DAPI (Akoya Biosciences). Slides were scanned using the VS120 Slide Scanning System (Olympus). Quantification was done using ImageJ software (National Institutes of Health) as described previously^91^).

### RNA purification from skin biopsies

Skin samples were cut into small pieces and suspended in 300 μl of lysis buffer (PureLink RNA Mini Kit – Thermo Fisher Scientific), prior to homogenisation on ice in 5-15 sec bursts for 2-5 min using IKA T10 homogeniser (T10 basic ULTRA-TURRAX). The homogenates were subsequently centrifuged at 16,000 x g for 5 minutes and the supernatants were transferred to a clean RNase-free tubes and total RNA extracted with the PureLink RNA Mini Kit (Thermo Fisher Scientific).

### Systemic R848 challenge

C57BL/6NCrl mice (used under Australian National University animal ethics, reference A2022/18) were injected intravenously with 200 μg of GGC-v1 conjugated with *in vivo*-jetPEI® (Polyplus #101000030) in 5% Glucose, 1 h prior to intraperitoneal injection with 25 μg R848 VacciGrade (InvivoGen #vac-r848). Two hours after injection of R848, mice were bled retroorbitally for collection of serum and sacrificed for spleen collection. The spleen was processed to a single-cell suspension and following lysis of red blood cells, total RNA was purified from 5×10^6^ total splenocytes using the Isolate II RNA mini-kit (Meridian Biosciences # BIO-52072) according to the manufacturer’s instructions. RNA was transcribed into cDNA using SuperScript IV Reverse Transcriptase (Thermo Fisher Scientific #18090010) according to the manufacturer’s instructions. RT-qPCR analyses were carried out with the Power SYBR Green Master Mix (Thermo Fisher Scientific) on an Applied Biosystems 7900 machine (Thermo Fisher Scientific). Each PCR was performed in technical triplicates with mouse *Gapdh* used as the reference gene. Relative gene expression was calculated using the 2^-ΔΔCt^ method. The primers used for Figure 5a were as follows: mIl10-FWD GCT CTT ACT GAC TGG CAT GAG, mIl10-REV CGC AGC TCT AGG AGC ATG TG; mTNF-FWD CGC TCT TCT GTC TAC TGA ACT TCG G, mTNF-REV AGAACTGATGAGAGGGAGGCCATTT; mIl6-FWD TCT ATA CCA CTT CAC AAG TCG GA mIl6-REV GAA TTG CCA TTG CAC AAC TCT TT; mGapdh-FWD AAT GTG TCC GTC GTG GAT, mGapdh-REV CTC AGA TGC CTG CTT CAC.

### Systemic FLuc mRNA administration

These animal experiments were conducted by Eurofins Discovery Pharmacology Discovery Services Taiwan, and approved by Institutional Animal Care and Use Committee reference IN020-08202020-27736. Female 8-week-old 129X1/SvJ mice (Jackson Laboratories) (∼25 g) were injected intravenously (i.v.) with ∼20 μg FLuc mRNA (actual 20.349 μg and 20.484 μg) encapsulated in LNPs with, or without, GGC-v1 (see details above). Bioluminescence imaging was performed at 6 h post-injection using an IVIS Spectrum®. Briefly, mice were anaesthetised with 4 % isoflurane, and 3 mg/mouse d-luciferin potassium salt in PBS was administered i.v. (in order to quantify luminescence expression using images recorded for 3 minutes, starting 5 minutes post-luciferin injection). For bioluminescence image analysis, regions of interest encompassing the area of signal were defined using the IVIS Spectrum, and the total number of photons per second [counts/second (cps)] was recorded. Blood was sampled at 6 h post-injection by submandibular bleed for serum quantification of IFN-α by ELISA (Invitrogen; BMS 6027 – analysed on TECAN Infinite F50 with Tecan i-control software) and other inflammatory cytokines by Bio-Plex Pro Mouse Cytokine 23-plex Assay (Biorad) on a BIO-RAD Bio-Plex 200 Luminex (using Bio-Plex Manager Software 6.0). Livers were also collected at 24 h and snap-frozen at -80°C until analysis. To measure luciferase activity, cell culture lysis reagent (CCLR, Promega) was added to fresh livers at 1 ml per 1 g of tissue, followed by homogenization with a polytron (<10 sec on ice). The samples were centrifuged at 16,000 g for 10 min at 4°C. Liver lysates were first normalized to 40 mg/mL with CCLR, then diluted 500X for luciferase activity measurement using a plate reader within 5 minutes. Luciferase activity was quantified as relative light units (RLU).

### Statistical analyses

Statistical analyses were carried out using Prism 10 (GraphPad Software Inc.). One-way and two-way analyses of variance (ANOVA) with uncorrected Fisher’s LSD were used when comparing groups of conditions, while unpaired two-tailed t-tests were used when comparing selected pairs of conditions. “ns” is non-significant.

## Acknowledgements

We thank K. Sakaniwa for the purified TLR8 protein; V. Hornung for the HEK-293T cells; C. David and Y. Crow for the TLR7 GOF expressing vectors; R. Veedu for advice on short oligonucleotide synthesis; E. Latz for the UNC93B1-mCitrine vector; M. Sweet for the RAW-ELAM cells; T. Wilson and L. J. Gearing for RNA sequencing and help with analyses; Eurofins Discovery for generating mRNA LNPs and conducting associated *in vivo* studies; B.R.G. Williams and S. Masters for comments on the manuscript; M. Kikkawa and Y. Sakamaki for managing and supporting the Graduate School of Medicine cryo-EM facility at the University of Tokyo. We acknowledge use of equipment and technical assistance of Monash Histology Platform, Department of Anatomy and Developmental Biology, Monash University; the Phenomics Translational Initiative teams for technical assistance with *in vivo* mouse studies including: K. Kwong and F-J. Li for the mouse intravenous and intraperitoneal injections and A. Davies and K. Diamand for cytokine and RT-qPCR measurements; the Sydney Analytical Core Research Facility (University of Sydney) for access to SPR infrastructure.

This work was supported by the Australian National Health and Medical Research Council Project Grant (2020565 to M.P.G., J.I.E and B.C.); mRNA Victoria Research Acceleration Fund, the Victorian Government’s Operational Infrastructure Support Program and COVID-19 Treatments Medical Research Fund; Grant-in-Aid from the Japanese Ministry of Education, Culture, Sports, Science, and Technology (Grant Nos. 22K15046 and 24K09349 to Z.Z., 22H02556 to U.O., 22H05184 and 23H00366 to T.S.); CREST, JST (Grant No. JPMJCR21E4 to T.S.); and Noxopharm Limited. Cryo-EM analyses were supported by the Basis for Supporting Innovative Drug Discovery and Life Science Research (BINDS) from the Japan Agency of Medical Research and Development (AMED) (Grand No. JP21am0101115; support No. 1570, 1846, 1848). This research project was undertaken with the assistance of resources and services from the National Computational Infrastructure and the Pawsey Supercomputing Research Centre, which are supported by the Australian Government and the Government of Western Australia accessed through the National Merit Allocation and ANU Merit Allocation schemes.

## Author contributions

Conceptualization: M.P.G, S.S., A.S.A., O.F.L., D.S.W., M.S., R.J., B.C., Z.Z., T.S.; Investigation: A.S.A., S.S., R.J., Z.Z., U.O., W.S.N.J., E.R., L.C., L.W.W., R.G., J.I.E., A.L.M., R.R., M.S., L.Y., H.H.F., J.B., S.H., D.Y., O.F.L., M.P.G.; Resources: K.A.L., P.H., C.G.V., M.A.B., U.O., O.F.L., D.S.W., M.S., B.C., T.S., M.P.G.; Data Curation: A.L.M., Z.Z.; Writing - Original Draft: M.P.G., B.C., Z.Z., U.O., T.S.; Writing - Review & Editing: S.S., M.S., O.F.L., R.J., D.S.W., E.R., W.S.N.J., S.H., J.I.E., H.H.F., L.W.W., A.L.M., D.Y., R.G., L.C., L.Y., Z.Z., U.O., C.G.V., K.A.L., M.A.B., T.S., B.C., M.P.G.; Supervision: M.P.G., B.C., T.S.; Project administration: A.S.A., S.S., Z.Z., U.O., R.J., M.S., O.F.L., B.C., T.S. and M.P.G.; Funding acquisition O.F.L., B.C., T.S., M.P.G.

## Data and materials availability

RNA sequencing data has been deposited in the NCBI Gene Expression Omnibus (GEO) with accession XXXX.

The cryo-EM maps have been deposited at the Electron Microscopy Data Bank under the following accession codes: EMD-60515 (TLR7/GUC-v1 complex) and EMD-60541 (TLR7/mGrArA complex). The coordinates of the atomic models have been deposited at the Protein Data Bank (PDB) under the following accession codes: TLR7/GUC-v1-*SS* (8ZW2), TLR7/GUC-v1-*RR* (8ZW4), TLR7/mGrArA-*SS* (8ZXE) and TLR7/mGrArA-*RR* (8ZXF).

---

**Supplementary Figure 1.**
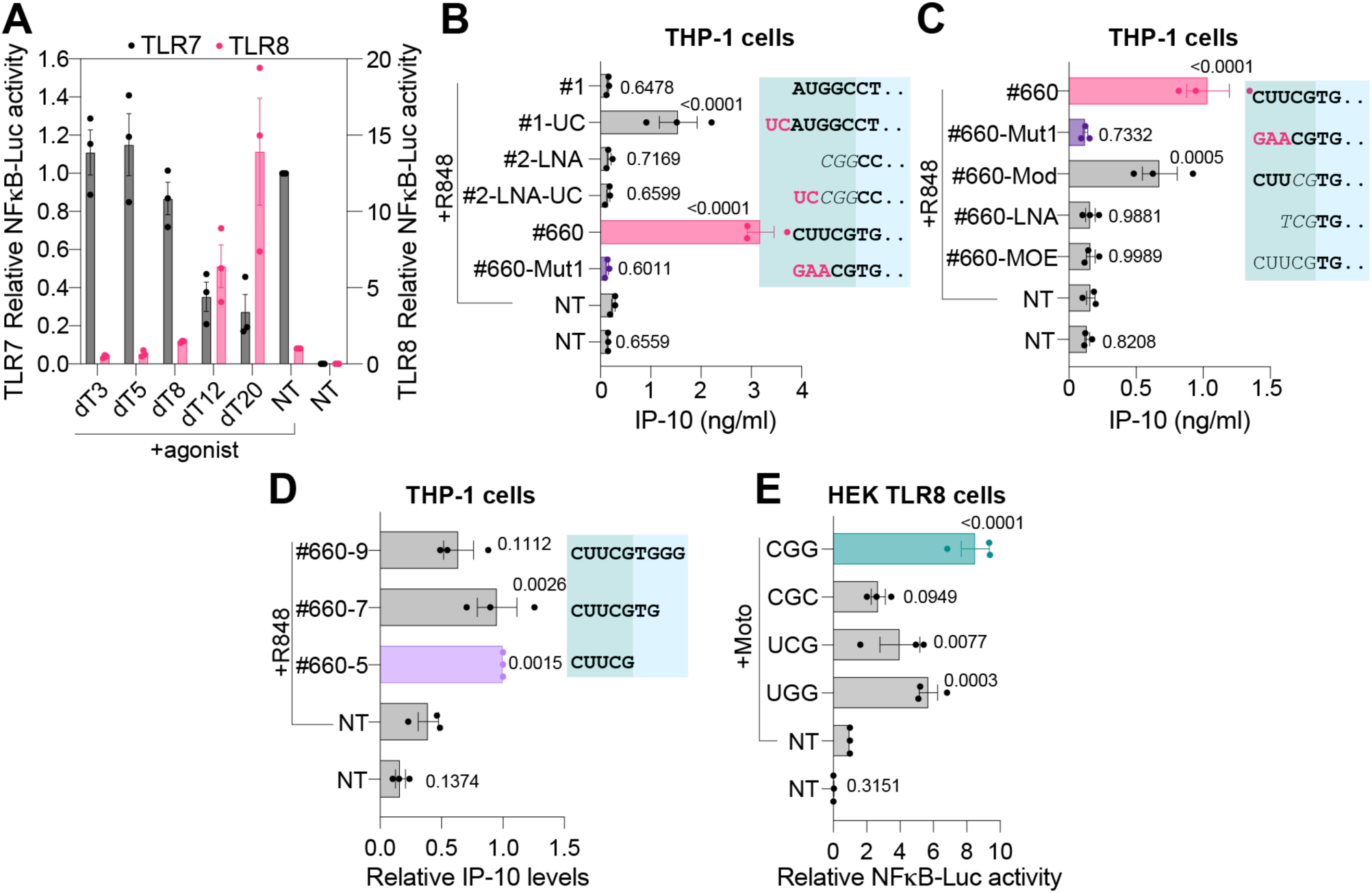
TLR8 potentiation by 3-mer oligos. (A) HEK TLR8 (right Y-axis) and HEK-TLR7 (left Y-axis) cells were pre-treated for ∼30 min with 1 μM dT series oligos prior to overnight stimulation with 1 μg/ml of R848 (TLR7) or 20 mM Uridine (TLR8) followed by luciferase assay. Data were background-corrected using the non-treated (NT) condition and are shown as expression relative to R848 or uridine only (± s.e.m). (B, C, D) Monocytic THP-1 cells were incubated overnight with 100 nM of oligo (B, C) or 1 μM of oligo (D) and stimulated with 1 μg/ml of R848 for 7 h prior to IP-10 ELISA analysis (± s.e.m. and one-way ANOVA with uncorrected Fisher’s LSD tests shown compared to the R848-only condition). (E) HEK TLR8 cells were pre-treated ∼30 min with 5 μM of the indicated oligos prior to overnight stimulation with 600 nM of motolimod (Moto) followed by luciferase assay. Data were background-corrected using the non-treated (NT) condition and are shown as relative expression to motolimod only (± s.e.m. and one-way ANOVA with uncorrected Fisher’s LSD tests shown compared to the Moto-only condition). (A-E) Data are shown as mean of n=3 independent experiments.

**Supplementary Figure 2.**
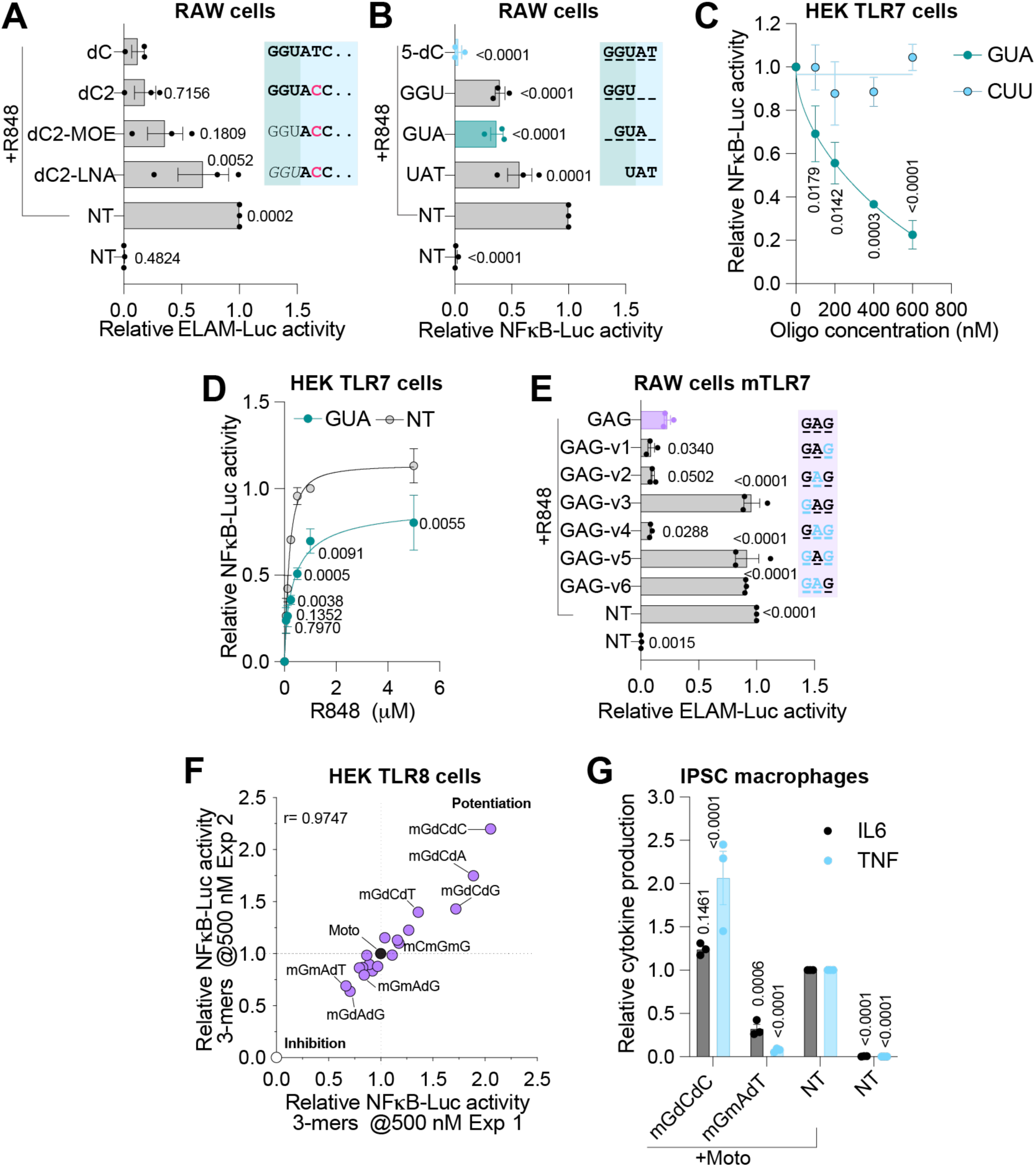
TLR7/8 modulation by 3-mer oligos. (A, B) Mouse RAW264.7 cells stably expressing an ELAM-Luciferase reporter (RAW-ELAM) were pre-treated for ∼60 min with 1 μM (A) or 5 μM (B) οf oligos, prior to overnight stimulation with 0.125 μg/ml R848 followed by luciferase assay. (C, D) HEK TLR7 cells were pre-treated for ∼30 min with the indicated concentration (0, 100, 200, 400 or 600 nM) (C) or 400 nM (D) of the oligos prior to overnight stimulation with 1 μg/ml (C) or indicated concentration (0, 0.062, 0.125, 0.25, 0.5, 1 μg/ml) of R848 followed by luciferase assay. (E) RAW-ELAM macrophages were pre-treated for 1 h with 5 μM GAG variants prior to overnight stimulation with 0.125 μg R848 followed by luciferase assay. (F) HEK TLR8 cells were pre-treated ∼60 min with 500 nM of the indicated oligos prior to overnight stimulation with 600 nM of motolimod (Moto) followed by luciferase assay. Pearson R correlation (r=0.9747 P<0.0001). (G) iPSC-derived macrophages were pre-treated for 30 min with 5 μM of oligos prior to stimulation with 400 nM motolimod (Moto) for 6 h and IL-6 and TNF ELISAs. Cytokine levels were normalised to the motolimod-only condition. Data were not corrected (D) or were background-corrected using the non-treated (NT) condition (A,B,C,E,F) and are shown as expression relative to the R848/motolimod-only conditions (± s.e.m. and one-way [A, B, E] or two-way [C, D, G] ANOVA with uncorrected Fisher’s LSD tests shown compared to dG [A], R848-only [B,D], _m_C_m_U_m_U [C], _m_G_m_A_m_G [E] or Moto [G] conditions). Data are mean of n=2 (D) or n=3 (A, B, C, E, G) independent experiments. (F) Data from two independent experiments averaged from 3 biological replicates are shown on each axis. (A, B) Green shading represents 5’-end base modification of the oligos as follows: bold is 2’-OMe, italic is LNA, non-bold is 2’-MOE. Black or pink DNA bases are shaded in light blue, and ”..” denotes that the sequence is truncated – see Table S5 for full-length sequences. (E) Black bases in bold denote 2’-OMe modification, and light blue bases denote DNA modifications. (F) _m_G_m_X_d_X and _m_G_d_X_d_X PS 3-mers were assessed.

**Supplementary Figure 3.**
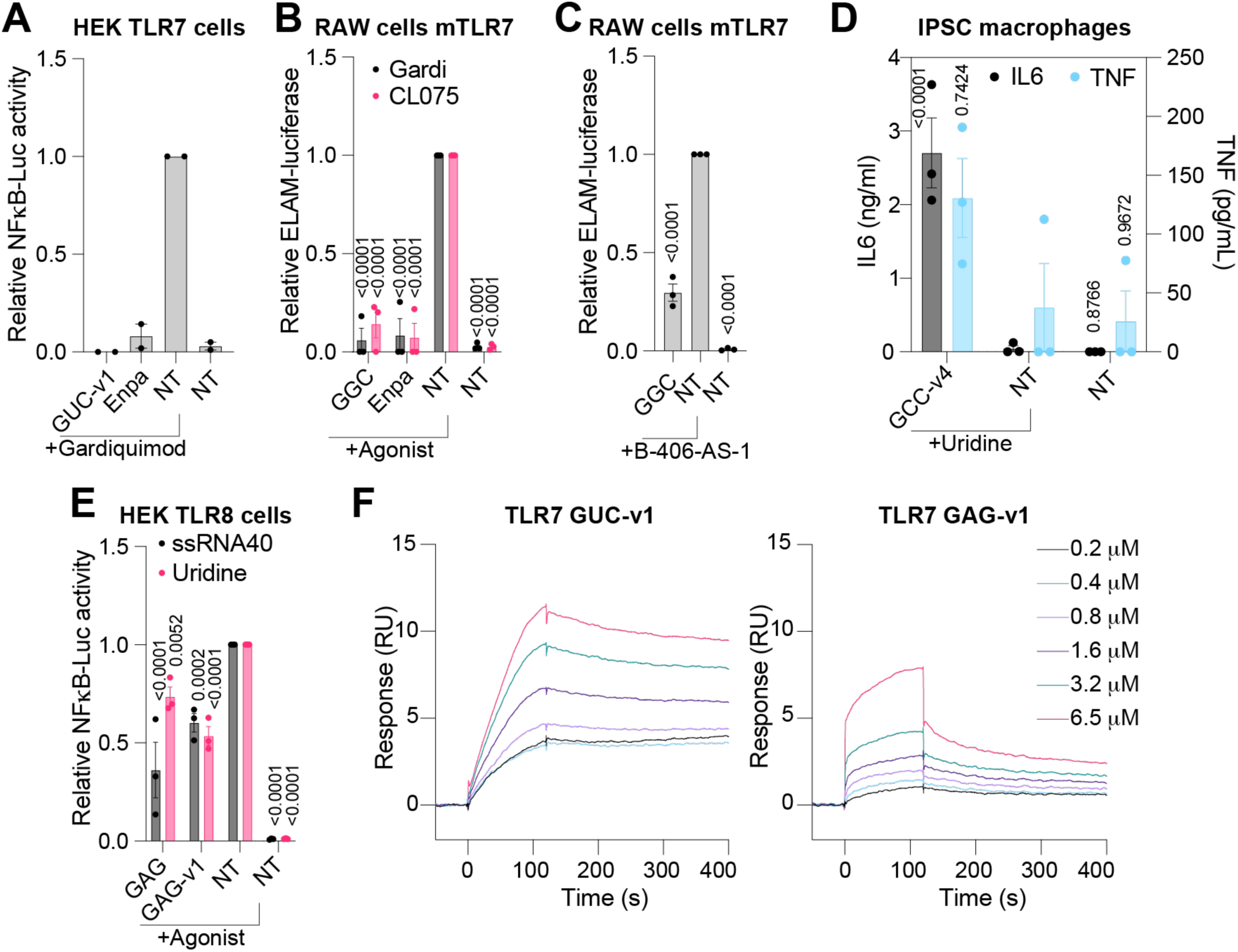
3-mer oligos bind to TLR7/8 to modulate their response to agonists. (A) HEK TLR7 cells were pre-treated for ∼30 min with 5 μM GUC-v1 or 50 nM enpatoran (Enpa) prior to overnight stimulation with 1 μg/ml of Gardiquimod followed by luciferase assay. (B, C) RAW-ELAM macrophages were pre-treated for ∼30 min with 5 μM (B) or 500 nM (C) _m_G_m_G_m_C prior to overnight stimulation with 0.5 μg/ml Gardiquimod or CL075 (B) or DOTAP transfection with 500 nM of B-406-AS-1 ssRNA (C) followed by luciferase assay. (D) iPSC-derived macrophages were pre-treated 30 min with 5 μM of GCC-v4 oligo prior to overnight stimulation with 20 mM uridine and IL-6 and TNF ELISAs. (E) HEK TLR8 cells were pre-treated for ∼120 min with 5 μM of _m_G_m_A_m_G or GAG-v1 prior to overnight stimulation with 20 mM of uridine or DOTAP transfection with 1 μM of ssRNA40 followed by luciferase assay. Data were not corrected (D) or were background-corrected (A-C,E) using the non-treated (NT) condition and are shown as expression relative to the agonist-only condition (± s.e.m. and one-way [C] or two-way ANOVA [B,D,E] with uncorrected Fisher’s LSD tests shown compared to agonist condition). Data are shown as mean of n=2 (A) or n=3 (B-E) independent experiments. (F) Surface plasmon resonance (SPR) analyses of recombinant *Macaca mulatta* TLR7 (with indicated concentrations of 3-mers). Data shown are representative of 5-6 independent analyses (Supplementary Table S2).

**Supplementary Figure 4.**
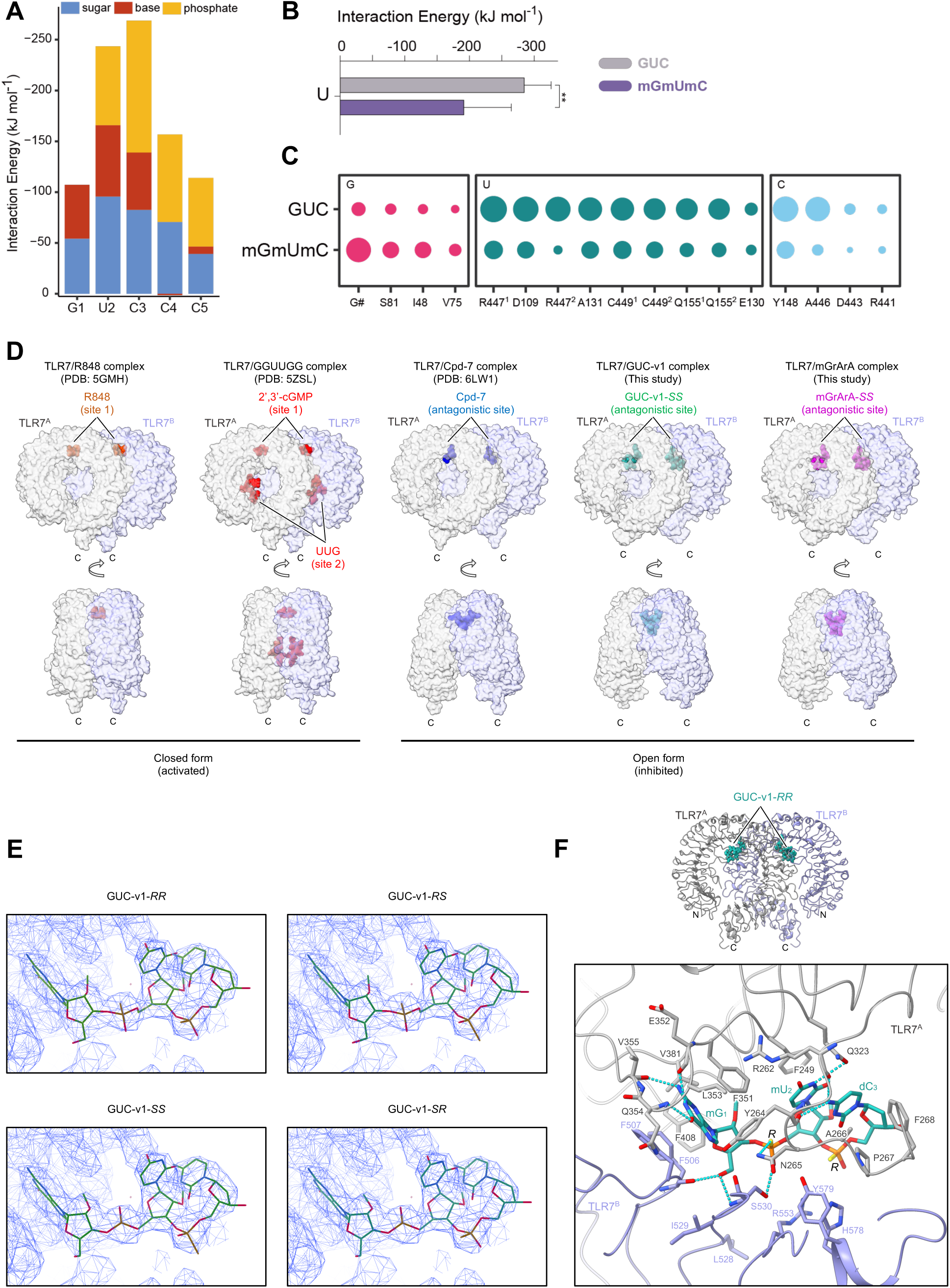
GUC-v1 binds at the antagonistic site of TLR7 leading to an open conformation. (A) The decomposed interaction energy of GUCCC with the protein residues that form binding site 2 from CphMD simulations at pH 5, suggesting the first three bases are essential for RNA binding and recognition. (B) The attractive interactions between the receptor and uridine are decreased by ∼60 kJ mol^-1^ with the uracil base leaving its binding site in _m_G_m_U_m_C simulations. (C) The overall percentage of interactions between the RNA bases and the protein residues that form binding site 2 retained during simulation are shown (the relative size of the circle represents the % time for observing each interaction pair formed along whole simulation trajectories). The intermolecular interactions between the nucleotides and receptor are decreased overall in _m_G_m_U_m_C simulations at site 2 compared with GUC. Labels on protein residues indicate different pairs of interactions between RNA ligands and residue side-chains. (D) Structural comparison of various TLR7/ligand complex structures. In each structure, the two TLR7 protomers TLR7A and TLR7B and ligands are colored by gray, purple and different colors, respectively. (E) Views of ligand fitting of four GUC-v1 stereoisomers into the cryo-EM map using COOT software. Cryo-EM map (level = 0.031) of the TLR7/GUC-v1 complex and the fitted GUC-v1 stereoisomers are shown in mesh and stick representations, respectively. (F) Overall structure of the TLR7/GUC-v1-RR complex (upper). Close-up view of GUC-v1-RR recognition at the antagonistic site.

**Supplementary Figure 5.**
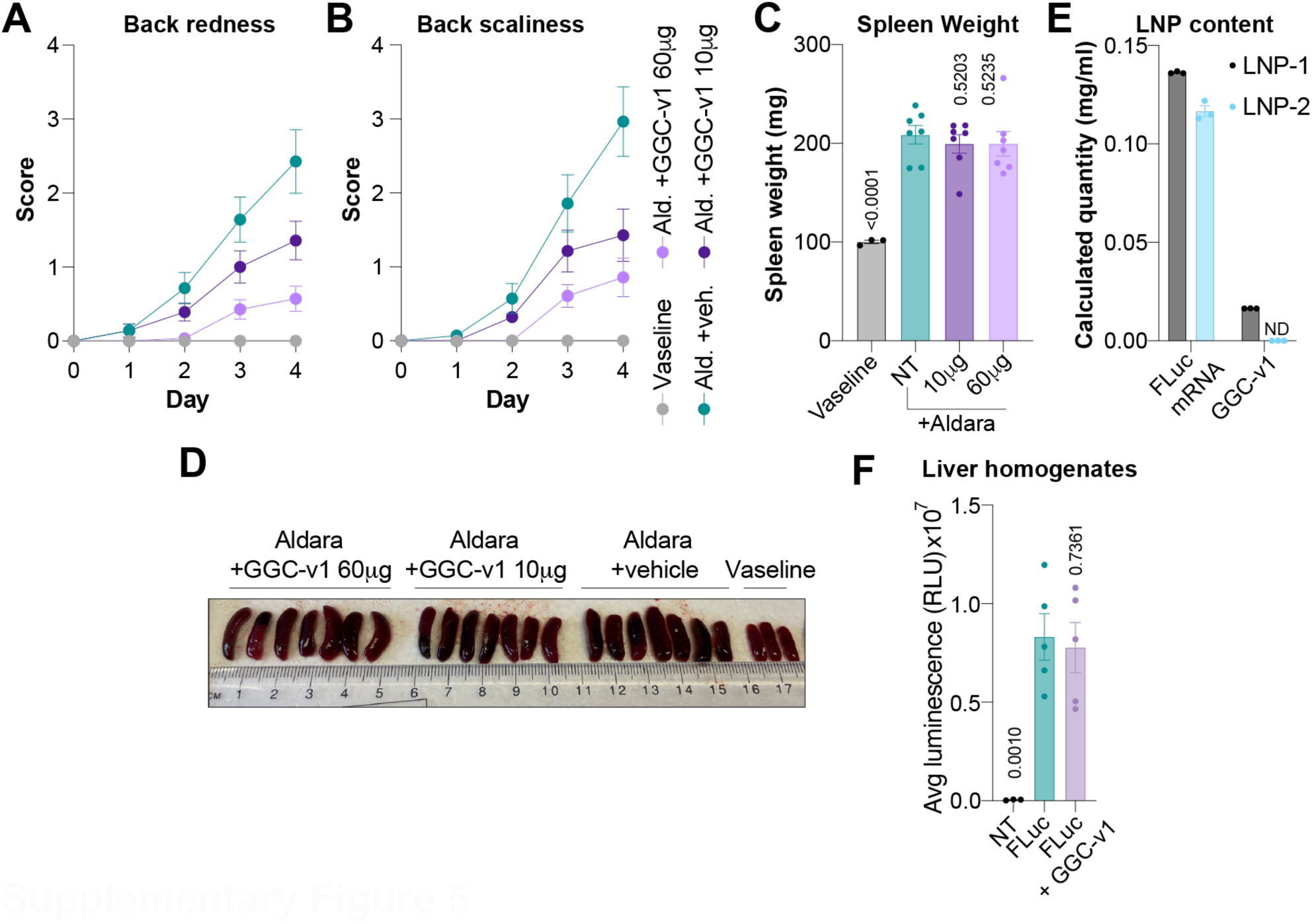
2’-OMe 3-mer oligos act as TLR7 antagonists *in vivo*. (A-C) WT C57/BL6 mice were treated with Aldara cream directly following, or not, application of 10 μg or 60 μg GGC-v1 formulated in F127 Pluronic gel for four days. Redness and scaliness were assessed by blinded investigators. After four days, mice were humanely euthanised and the spleens collected, weighed (C), and photographed (D). (C) Mean of n = 3-7 mice/group is shown (± s.e.m. and one-way ANOVA with uncorrected Fisher’s LSD tests shown compared to Aldara-only group). (E) RT-qPCR of FLuc mRNA and LC-MS quantification of GGC-v1 from two LNP preparations – both are absolute quantification with the standard curve, FLuc mRNA, or GGC-v1. ND: Not Detected at limit level of detection (10 ng/mL). LNP1 contains FLuc+GGCv1 and LNP2 contains FLuc mRNA only. (F) Luciferase activity measured from liver homogenates at 24 h post injection. Mean of n = 3-5 mice/group is shown (± s.e.m. and one-way ANOVA with uncorrected Fisher’s LSD tests shown compared to FLuc mRNA-only group).

**Supplementary Figure 6.**
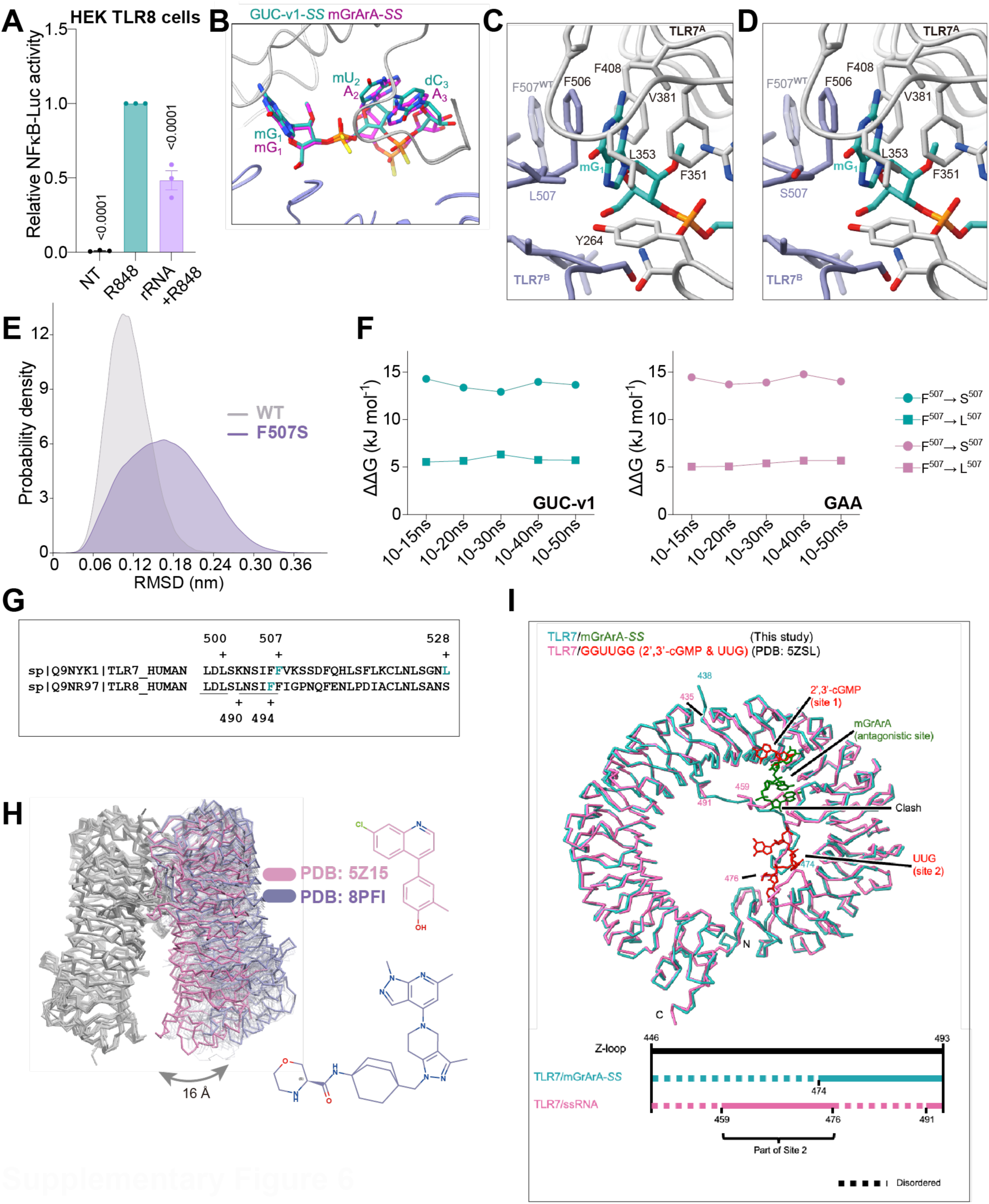
Endogenous 2’-OMe RNA binds to a conserved TLR7/8 antagonistic pocket. (A) HEK TLR8 cells were transfected for 6 h with 3 μg/ml purified rRNA prior to overnight stimulation with 1 μg/ml of R848 followed by luciferase assay. (B) Comparison of the conformations of GUC-v1-*SS* and _m_G_r_A_r_A-*SS* at the TLR7 antagonistic site. Structure of PO-GUC-v1 at the antagonistic binding site of mutant TLR7 F507L (C) and TLR7 F507S (D) from MD simulations. The phenylalanine at position 507 in wild-type TLR7 is shown in transparent licorice. Carbon atoms in different protomers are colored in grey and steel blue, carbon atoms in GUC-v1 are colored in cyan, O, N, and P atoms are in red, blue, and orange, respectively. Protein ribbons are in grey and steel blue separately. (E) The distribution of RMSD value of _m_G_1_ of GUC-v1 in wild-type (grey) and F507S mutant (steel blue) in an MD simulation at a constant pH of 5. (F) Convergence of the relative binding free energy difference of the antagonistic 3-mers using different length of trajectories in free energy perturbation calculations. (G) Protein sequence alignments of human TLR7 and TLR8 antagonistic regions – the residues underlined are entirely conserved. TLR7 F507, L528, and TLR8 F494 are highlighted in blue. (H) The different conformations of an antagonist binding to inactive TLR8 dimers superimposed using single protomer chain with structures from PDB 5WYX, 5WYZ, 5Z14, 5Z15, 6KYA, 6TY5, 6V9U, 6ZJZ, 7CRF, 7R52, 7R53, 7R54, 7RC9, 7YTX, and 8PFI. The structures of inhibitory molecules bound in the most compact inactive dimer (5Z15, pink) and the most open inactive dimer (8PFI, steel blue) are shown on the right, with _m_G_r_A_r_A docking best to the most open dimer. (I) Structural comparison of the protomer structures from the TLR7_/m_G_r_A_r_A-*SS* complex and the TLR7/GGUUGG complex (PDB: 5ZSL). One TLR7 protomer in each dimer structure was aligned using the ChimeraX matchmaker tool and is shown in main chain trace. The ordered and disordered regions of Z-loop in each structure are indicated at the bottom. (A) Data were background-corrected using the NT condition and are shown as expression relative to the R848-only condition (± s.e.m. and one-way ANOVA with uncorrected Fisher’s LSD tests shown compared to R848-only condition [A]). (A) Data are shown as the mean of n=3 independent experiments.

**Supplementary Figure 7.**
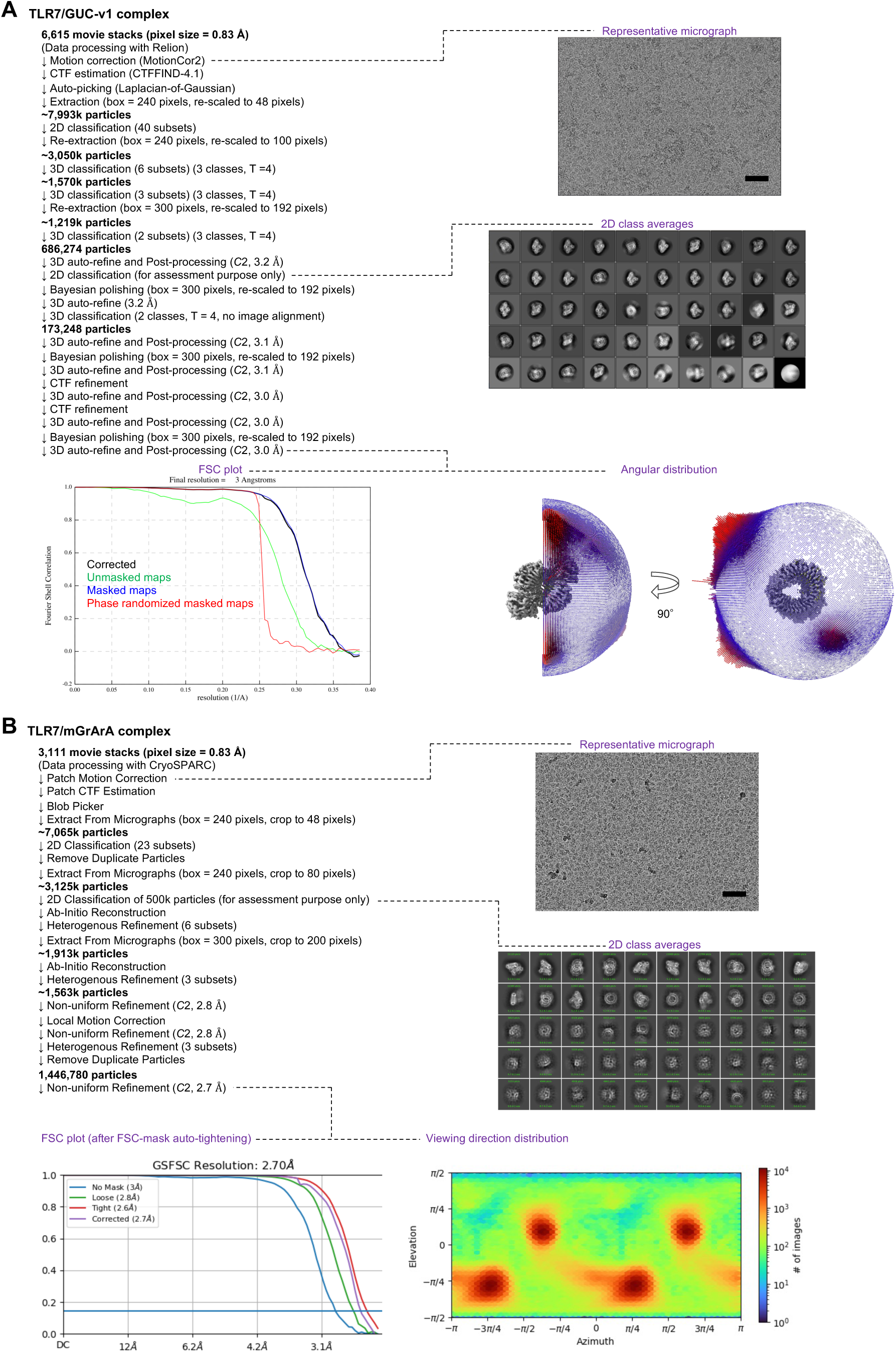
Cryo-EM data processing. Data processing workflow of cryo-EM analysis of TLR7/GUC-v1 complex (A) and TLR7/mGrArA complex (B). Representative motion-corrected micrographs (out of 6,615 (A) and 3,111 (B) total micrographs), 2D class averages, Fourier shell correlation (FSC) plot of the final 3D reconstruction (resolution cut-off at FSC = 0.143), and the angular distribution (A) or viewing direction distribution (B) are shown.

**Supplementary Table S1.**
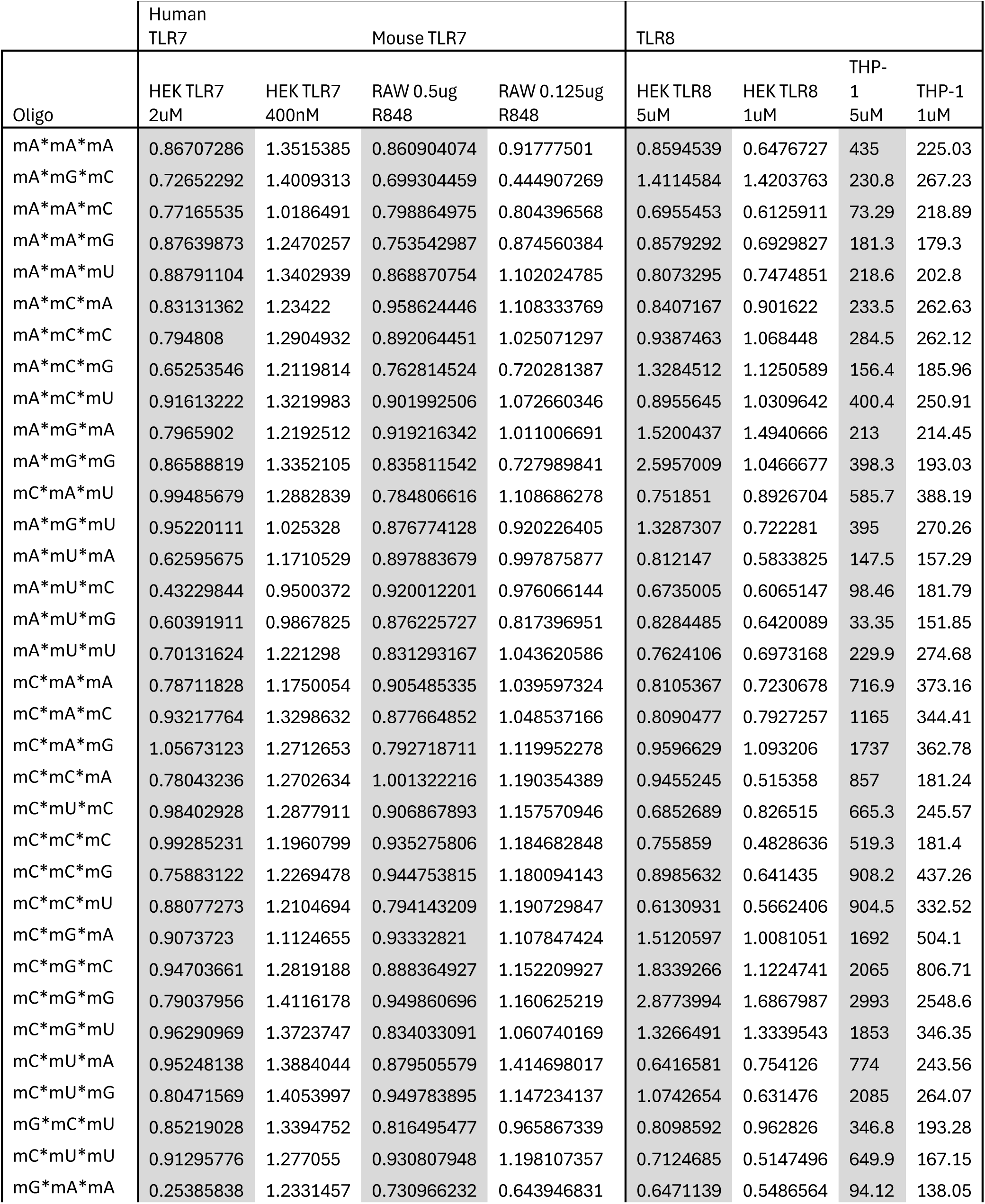
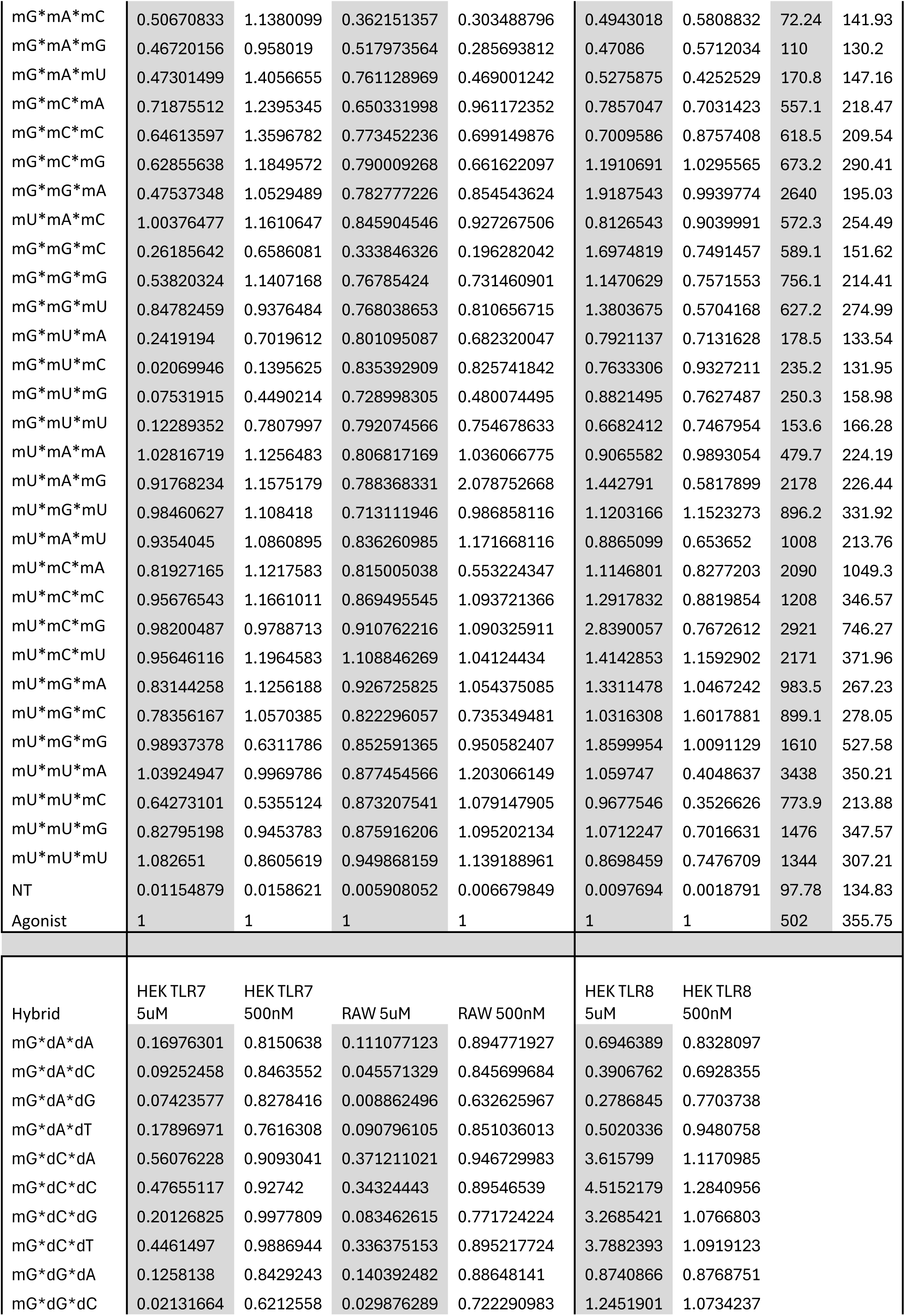
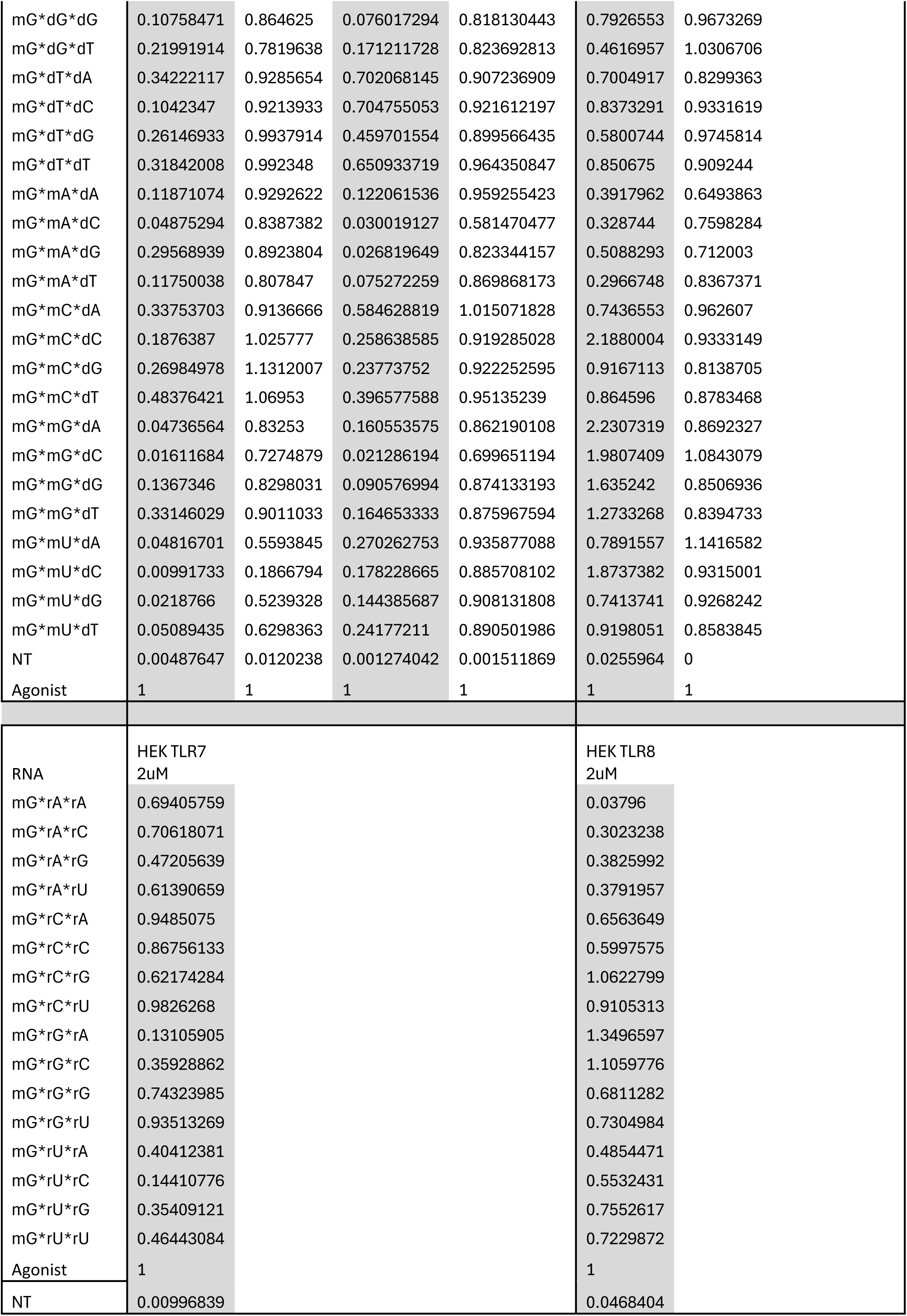
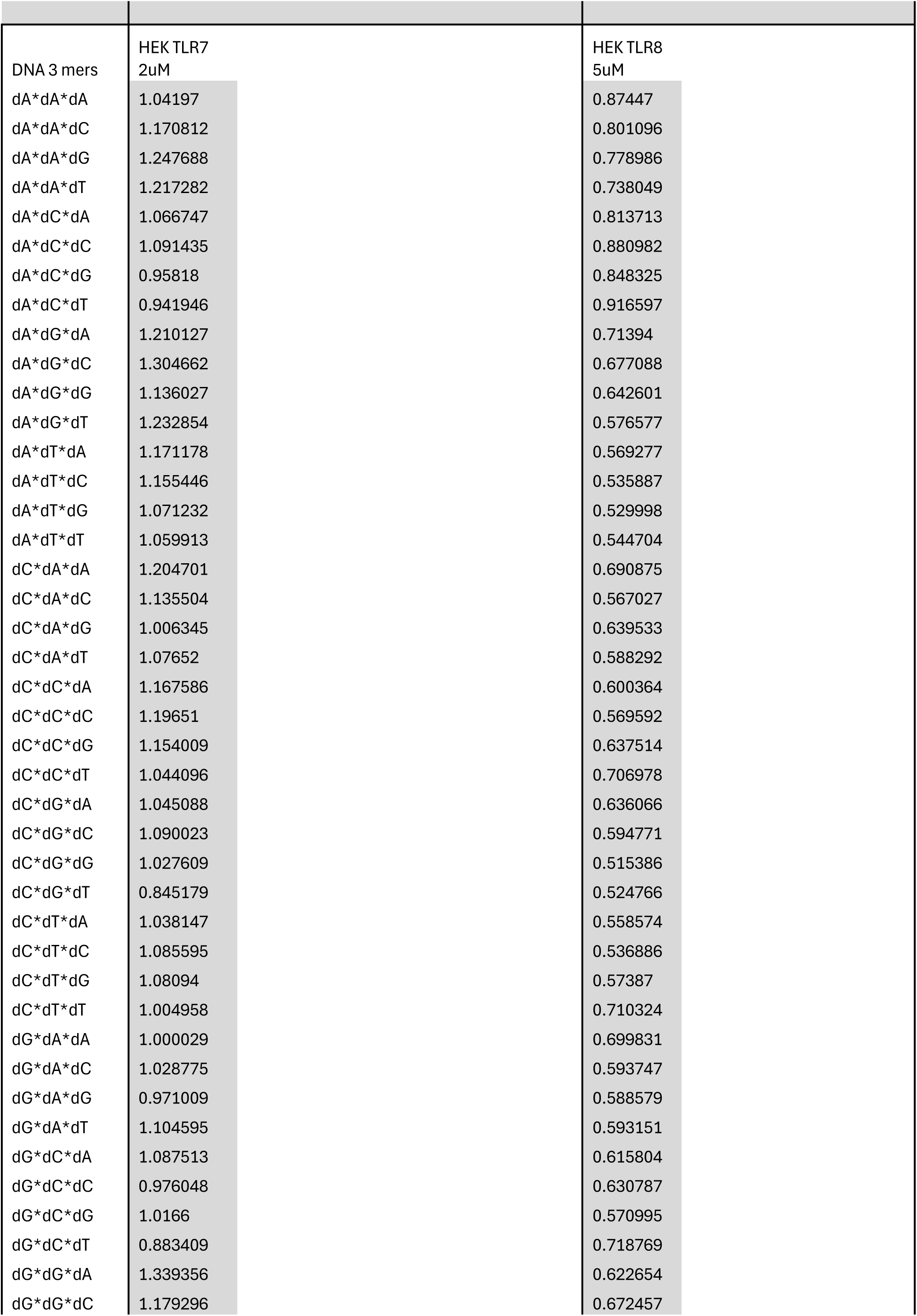
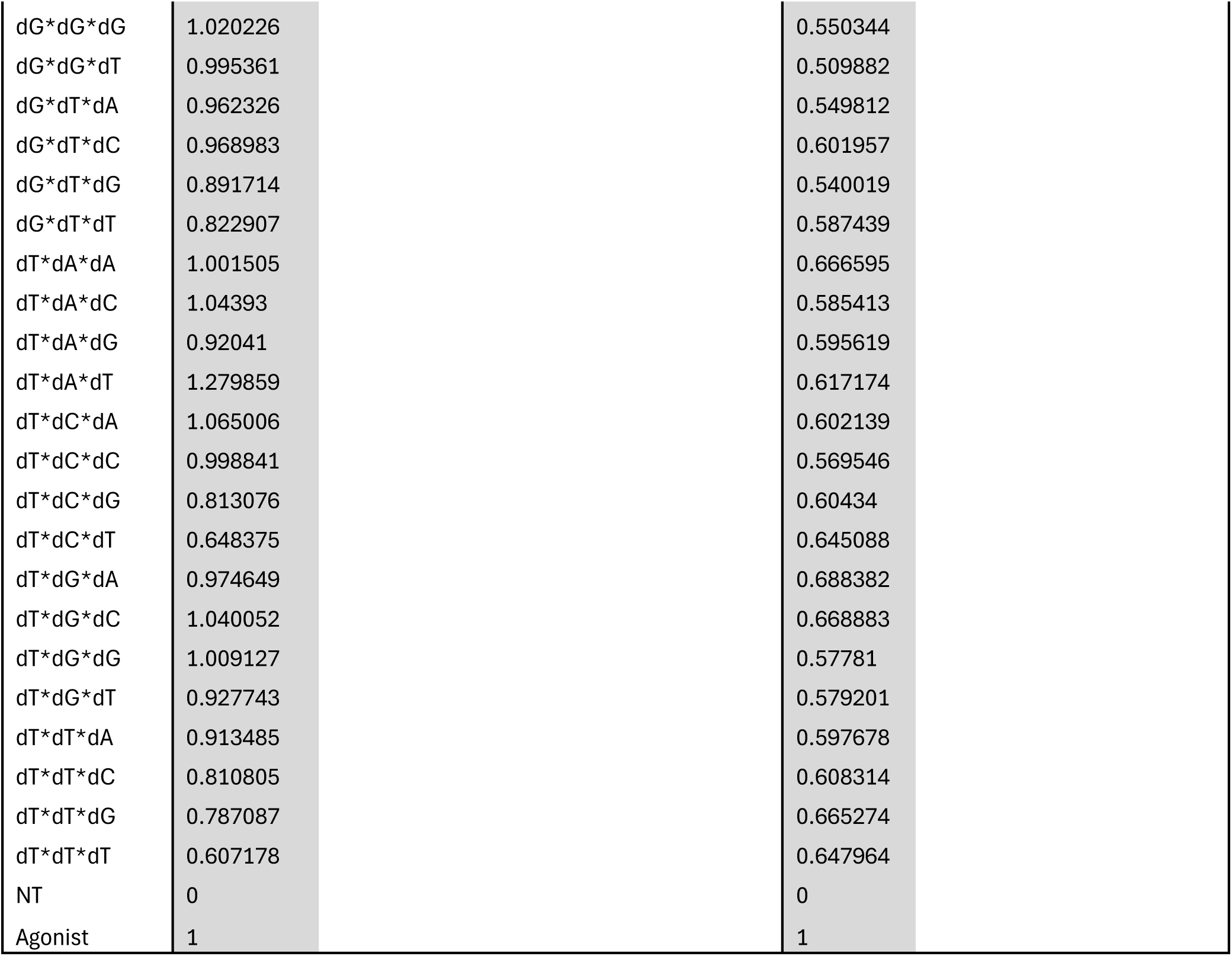
Oligonucleotides screen data. 2′-OMe is mX, DNA is dX, RNA is rX, and phosphorothioate inter-nucleotide linkages are denoted with a *. All values were background corrected and reported to agonist only condition, except for the screens in THP-1 cells were raw IP-10 concentrations (pg/ml) are provided. The concentrations provided indicate the dose of the 3-mers used in the screens except for the 2’-OMe 3-mer screens in mouse RAW-ELAM cells were the dose provided (μg/ml) are that of R848 stimulation (the oligos were at 5 μM).

**Supplementary Table S2:** SPR analyses.

(A) Affinity (KD) values for SPR runs with TLR7 and oligonucleotides.
|  | Affinity Values (μM) |  |  |  |  |
| --- | --- | --- | --- | --- | --- |
|  | Run 1 | Run 2 | Run 3 | Run 4 | Run 5 |
| GUC_V1 | 7 ± 2 | 7 ± 1 | 6 ± 1 | 4 ± 2 | 4 ± 2 |
| GAG_V1 | 38 ± 2 | 32 ± 2 | 30 ± 1 | 20 ± 11 | 11 ± 1 |
| GUC-V6 | Negligible binding observed |  |  |  |  |
| GCC-V4 | Negligible binding observed |  |  |  |  |

(B) Affinity (KD) values for SPR runs with TLR8 and oligonucleotides.
|  | Affinity Values (μM) |  |  |  |  |  |
| --- | --- | --- | --- | --- | --- | --- |
|  | Run 1 | Run 2 | Run 3 | Run 4 | Run 5 | Run 6 |
| GAG_V1 | 3 ± 1 | 4 ± 1 | 5 ± 1 | 3 ± 0.4 | 4 ± 1 | 4 ± 1 |
| GCC-V4 | 5 ± 1 | 7 ± 1 | 5 ± 1 | 12 ± 3 | 7 ± 1 | 11 ± 2 |
| GUC-V6 | Negligible binding observed |  |  |  |  |  |
| GUC_V1 | Not detected |  |  |  |  |  |

**Supplementary Table S3.** Cryo-EM data collection, refinement and validation statistics.

| Data | TLR7/GUC-v1 | TLR7/mGrArA |
| --- | --- | --- |
| **EMDB ID** | EMD-60515 | EMD-60541 |
| **PDB ID** | PDB 8ZW2 (GUC-v1-SS)<br>PDB 8ZW4 (GUC-v1-RR) | PDB 8ZXE (mGrArA-SS)<br>PDB 8ZXF (mGrArA-RR) |
| **Data collection and processing** |  |  |
| Microscope | Titan Krios G4 | Titan Krios G4 |
| Voltage (kV) | 300 | 300 |
| Detector | K3 | K3 |
| Magnification | 105,000 | 105,000 |
| Pixel size (Å) | 0.83 | 0.83 |
| Total dose (e<sup>-</sup>/Å<sup>2</sup>) / Frames per movie | ~48 / 48 | ~48 / 48 |
| Total movie stacks | 6,615 | 3,111 |
| Software | RELION v4.0.1 | cryoSPARC v4.0.3 |
| Final particle images (no.) | 173,248 | 1,446,780 |
| Symmetry imposed | C2 | C2 |
| Map resolution (Å) | 3.0 | 2.7 |
| FSC threshold | 0.143 | 0.143 |
| **Refinement** |  |  |
| Software | Chimera, COOT, JLigand, Phenix | Chimera, COOT, JLigand, Phenix |
| Model resolution (Å)<br>(FSC threshold = 0.5) | 3.4 / 3.5 | 3.3 / 3.3 |
| Model composition |  |  |
| Protein chains | 2 / 2 | 2 / 2 |
| Ligands | 2×GUC-v1-SS<br>or GUC-v1-RR | 2×mGrArA-SS<br>or mGrArA-RR |
| Residues | 1534 / 1534 | 1526 / 1526 |
| Average B-factors (Å<sup>2</sup>) | 127.2 / 131.9 | 124.6 / 124.6 |
| R.m.s deviations |  |  |
| Bond lengths (Å) | 0.003 / 0.003 | 0.005 / 0.005 |
| Bond angles (°) | 0.540 / 0.514 | 1.015 / 1.015 |
| Validation |  |  |
| Molprobity score | 1.74 / 1.86 | 1.64 / 1.64 |
| Clashscore | 5.68 / 5.68 | 5.32 / 5.32 |
| Poor rotamers (%) | 1.0 / 1.4 | 0.7 / 0.7 |
| Ramachandran plot |  |  |
| Favored (%) | 93.5 / 93.0 | 94.9 / 94.9 |
| Allowed (%) | 6.1 / 6.6 | 4.8 / 4.8 |
| Outliers (%) | 0.4 / 0.4 | 0.4 / 0.4 |

**Supplementary Table S4.** Fitted pKa values of all titratable residues (D: Aspartate, E: Glutamate, H: Histidine) using PO-GUC-v1 binding inactive TLR7 dimer structure from constant pH MD titration experiments with a range of pH from 2.0 to 10.5.

|  | resid | pka |
| --- | --- | --- |
| D | 37 | <2 |
| D | 41 | 4.6 |
| D | 50 | 2.9 |
| D | 53 | 3.2 |
| D | 79 | 4.5 |
| D | 89 | 4.6 |
| D | 95 | 5.2 |
| D | 135 | <2 |
| D | 198 | 5 |
| D | 213 | 6.1 |
| D | 243 | 4.2 |
| D | 244 | 5.3 |
| D | 255 | <2 |
| D | 287 | <2 |
| D | 320 | <2 |
| D | 332 | 2.3 |
| D | 346 | 2.2 |
| D | 403 | <2 |
| D | 427 | <2 |
| D | 469 | 3.6 |
| D | 499 | >10.5 |
| D | 512 | <2 |
| D | 548 | <2 |
| D | 555 | >10.5 |
| D | 572 | 5.7 |
| D | 605 | 4.2 |
| D | 607 | 5.9 |
| D | 632 | 2.8 |
| D | 639 | 2.3 |
| D | 656 | <2 |
| D | 671 | 3.4 |
| D | 705 | 5.5 |
| D | 745 | 4.6 |
| D | 753 | <2 |
| D | 790 | 2.8 |
| D | 811 | 2.9 |
| D | 829 | 4.8 |
| E | 58 | 4 |
| E | 93 | <2 |
| E | 141 | 3.7 |
| E | 156 | 3.5 |
| E | 166 | 3.9 |
| E | 170 | 8 |
| E | 175 | 4.5 |
| E | 196 | <2 |
| E | 229 | 4.5 |
| E | 242 | 3.8 |
| E | 291 | 3.9 |
| E | 318 | 4.3 |
| E | 329 | <2 |
| E | 352 | <2 |
| E | 384 | 3.2 |
| E | 400 | 3.4 |
| E | 461 | <2 |
| E | 537 | 10.1 |
| E | 543 | 3.1 |
| E | 563 | 3.6 |
| E | 564 | 6.5 |
| E | 569 | 10.4 |
| E | 583 | 2.3 |
| E | 617 | 3.5 |
| E | 619 | 2.6 |
| E | 625 | 3.6 |
| E | 637 | 5.1 |
| E | 653 | 4.2 |
| E | 654 | 2.6 |
| E | 702 | 5.1 |
| E | 716 | 3 |
| E | 769 | 3.4 |
| E | 802 | 4 |
| H | 46 | 4.6 |
| H | 55 | 8 |
| H | 76 | 6 |
| H | 86 | 5.5 |
| H | 90 | 6.2 |
| H | 298 | 6.7 |
| H | 304 | 7.1 |
| H | 337 | 5.7 |
| H | 394 | 5.3 |
| H | 464 | 5.3 |
| H | 515 | 4.2 |
| H | 558 | 6.3 |
| H | 566 | 6.9 |
| H | 578 | 7.1 |
| H | 587 | 8.6 |
| H | 630 | 5.1 |
| H | 708 | 3 |
| H | 781 | 6.5 |
| H | 782 | 7.4 |
| H | 800 | 5.4 |

**Supplementary Table S5.** Oligonucleotide sequences. 2’-MOE is moX, 2′-OMe is mX, DNA is dX, RNA is rX, LNA is lX, and phosphorothioate inter-nucleotide linkages are denoted with a *.

|  |  |
| --- | --- |
| ASO #1 | mA*mU*mG*mG*mC*C*T*T*T*C*C*G*T*G*C*mC*mA*mA*mG*mG |
| ASO #1-UC | mU*mC*mA*mU*mG*mG*mC*C*T*T*T*C*C*G*T*G*C*mC*mA*mA*mG*mG |
| ASO #2-LNA | lC*lG*lG*C*C*T*C*G*G*A*A*G*C*lT*lC*lT |
| ASO #2-LNA-UC | mU*mC*lC*lG*lG*C*C*T*C*G*G*A*A*G*C*lT*lC*lT |
| ASO #660 | mC*mU*mU*mC*mG*T*G*G*G*G*T*C*C*T*T*mU*mU*mC*mA*mC |
| ASO #660-Mut1 | mG*mA*mA*mC*mG*T*G*G*G*G*T*C*C*T*T*mU*mU*mC*mA*mC |
| ASO #660-Mod | mC*mU*mU*lC*lG*T*G*G*G*G*T*C*C*T*T*mU*mU*mC*mA*mC |
| ASO #660-LNA | lT*lC*lG*T*G*G*G*G*T*C*C*T*T*lT*lT*lC |
| ASO #660-MOE | moC*moT*moT*moC*moG*T*G*G*G*G*T*C*C*T*T*moT*moT*moC*moA*moC |
| #660-5 | mC*mU*mU*mC*mG |
| #660-7 | mC*mU*mU*mC*mG*T*G |
| #660-9 | mC*mU*mU*mC*mG*T*G*G*G |
| dC | mG*mG*mU*A*T*C*C*C*C*C*C*C*C*C*C*C*C*C |
| dC2 | mG*mG*mU*A*C*C*C*C*C*C*C*C*C*C*C*C*C*C*C |
| dC2-MOE | moG*moG*moT*dA*dC*dC*dC*dC*dC*dC*dC*dC*dC*dC*dC*dC*dC |
| dC2-LNA | lG*lG*lT*dA*dC*dC*dC*dC*dC*dC*dC*dC*dC*dC*dC*dC*dC |
| 5-dC | mG*mG*mU*dA*dT |
| ssRNA40 | rG*rC*rC*rC*rG*rU*rC*rU*rG*rU*rU*rG*rU*rG*rU*rG*rA*rC*rU*rC |
| B-406-AS1 | rUrArArUrUrGrGrCrGrUrCrUrGrGrCrCrUrUrCrUrU |

